# PD-L1 – PD-1 interactions limit effector Treg cell populations at homeostasis and during infection

**DOI:** 10.1101/2020.12.09.416990

**Authors:** J.A. Perry, J.T. Clark, J. Gullicksrud, J. DeLong, L. Shallberg, B. Douglas, A. Hart, C. Konradt, J. Park, A. Glatman-Zaretzky, R. de Waal Malefyt, D.A. Christian, A. H. Sharpe, C.A. Hunter

## Abstract

While much is known about the factors that promote the development of diverse Treg cell responses, less is known about the pathways that constrain Treg cell activities. The studies presented here reveal that at homeostasis there is a population of effector Treg cells that express PD-1, and that blockade of PD-L1 or loss of PD-1 results in increased Treg cell activity. In response to infection with the parasite *T. gondii*, the early production of IFN-γ results in widespread upregulation of PD-L1. Moreover, blockade of PD-L1, whole body deletion of PD-1, or lineage-specific deletion of PD-1 in Foxp3^+^ cells prevented the loss of the effector Treg cells but resulted in reduced pathogen specific CD4^+^ T cell responses during infection. Thus, at homeostasis basal PD-L1 expression constrains and tunes the pool of Treg cells, but during infection the upregulation of PD-L1 provides a mechanism to contract the Treg cell population required to maximize the development of pathogen specific CD4^+^ T cell responses.

## Introduction

T cells are essential for surveillance and protection against cancer and infectious diseases, but there are multiple mechanisms that limit effector T cell responses in order to prevent auto-immunity and immune mediated collateral damage^1–3^. It is now established that Treg cells are a critical component of the regulatory mechanisms that limit aberrant T cell responses, and the loss of Treg cells is associated with a variety of immune mediated conditions^4–6^. Consequently, there has been a long-standing interest in defining the cellular and molecular pathways that promote Treg cell development and maintenance^7–9^. For example, Foxp3 is the lineage defining transcription factor required for the generation and maintenance of suppressive function in Tregs^6,10,11^. Likewise, at homeostasis Treg cells are activated by constitutively expressed self-antigen^12^, ongoing TCR signaling^13^ and costimulation^14,15^, and these processes contribute to the maintenance of the Treg cell pool. The ability of Treg cells to limit immune activity is mediated through numerous suppressive mechanisms that include the consumption of IL-2^16^, the secretion of IL-10^17^ and the expression of inhibitory ligands such as PD-L1, CTLA-4, and LAG3^18–20^.

Consistent with the presence of multiple suppressive mechanisms it is now appreciated that there are subsets of Treg cells that are associated with different classes of inflammation^21–23^. For example, Treg cells exposed to the cytokines IFN-γ, IL-12, or IL-27 upregulate the Th1 associated transcription factor T-bet, express CXCR3, and during infection these Th1-Treg cells are specialized to operate at sites of Th1-mediated inflammation^24–26^. More recently, effector Treg cells (eTregs) have been identified as Treg cells that undergo high affinity TCR-antigen interactions associated with increased expression of IL-10 and inhibitory receptors (IRs), while Treg cells with low-affinity TCR interactions have increased levels of CD25^27–29^. As such, there is evidence that these eTreg cells express the inhibitory receptor (IR) PD-1^30–33^, and elevated levels of TIGIT, GITR, PD-L1, and CTLA-4 which contribute to the suppressive functions of the eTreg^22,27,29,34^. For effector T cells, expression of PD-1 is associated with repeated TCR stimulation^32,35,36^ and the ability of this inhibitory receptor to antagonize costimulatory and TCR mediated signals^37,38^ culminates in the loss of function and eventual deletion of the T cell^39,40^. This pathway is also relevant to Treg cell biology, and their expression of PD-L1 allows them to limit T cells that express PD-1^41,42^. However, although PD-1 is not essential for thymically or peripherally derived Treg cells^42,43^, other reports concluded that activation of PD-1 supported the formation of inducible (i)Treg cells^34,41,44,45^. Moreover, recent studies have found that that blockade of the PD-1 pathway for cancer treatment results in increased numbers of PD-1^hi^ intra-tumor eTreg cells, associated with immune suppression, metastasis, and increased morbidity^32,33,45,46^.

During infection, the role of Treg cells is complex and increased Treg cell activity can favor pathogen persistence^47–49^, while infection can lead to the emergence of specialized subsets of Treg cells that limit immune pathology^24–26^. Paradoxically, many systemic pathogens result in an acute collapse of Treg cells and these events allow the emergence of effector responses required to control infection^47,50–52^. The basis for this phenomenon remains unclear but many infections lead to a suppression of basal IL-2 production^8,51,53–55^, a cytokine required for the maintenance of Treg cells^47^. Indeed, treatment of infected mice with IL-2 complexes that target CD25 mitigate this loss of Treg cells ^47,51,56^[HCA1][JP2]. The data presented here indicate that at homeostasis basal expression of PD-L1 restricts a population of PD-1^+^ eTreg cells but in response to infection with *T. gondii*, the production of IFN-γ results in marked upregulation of PD-L1 and contraction of the PD-1^+^ eTreg cell populations. However, blockade of PD-L1 or loss of PD-1 on Treg cells resulted in reduced parasite specific CD4^+^ effector T cells and mitigates the development of immune pathology. Thus, Treg cell expression of PD-1 provides a mechanism to rapidly tune eTreg cell activity in response to inflammatory signals.

## Materials and Methods

### Mice

C57BL/6 mice were purchased from Taconic (Rensselaer, NY, USA) at 6 weeks of age and housed in the University of Pennsylvania Department of Pathobiology vivarium in accordance with institutional guidelines. PD-1^−/−^ mice bred on the C57BL/6NTac background were created by deleting exons 2 and 3 from the *Pdcd1* locus through use of CRISPR/Cas9 gene editing at Taconic Artemis^57^. These PD-1^−/−^ mice were then bred and maintained by Taconic Biosciences Inc on behalf of Merck & Co., Inc. (Kenilworth, NJ, USA) and shipped to the University of Pennsylvania. Nur77^GFP^ mice on the C57BL/6J background created in the Hogquist Lab^58^ at the University of Minnesota were bred by The Jackson Laboratory (Bar Harbor, ME, USA) and shipped to the University of Pennsylvania. Foxp3^cre^ x PD-1^flox^ mice, generated by the Sharpe Lab^59^ at Harvard Medical School were shipped and then bred in the University of Pennsylvania vivarium in accordance with institutional guidelines. Foxp3^cre^ x PD-1^flox^ mice generated at the University of Pennsylvania were PCR screened using digested tail tips, with Foxp3^cre^ forward primer sequence of: 5’ - AGG ATG TGA GGG ACT ACC TCC TGT A -3’, and reverse primer sequence of: 5’ - TCC TTC ACT CTG ATT CTG GCA ATT T - 3’, under the condition settings of 3min at 94C, with 30 cycles of 45sec at 94C for melting temp, 30sec at 55C for annealing, and 1min at 72 sec for extension with a 2 minute final extension at 72C. For PD-1^flox^ forward primer sequence of: 5’ - GTC TCA ACA GAG GCC AGA GG - 3’ and reverse sequence of TGA AGG CTC CTC CTT CTT CA - 3’, under the condition settings of 4min at 94C, with 30 cycles of 45sec at 94C for melting temp, 30sec at 55C for annealing, and 1 min at 72C for extension, with a 10 minute final extension at 72C.

### Infection and Blockade

Infections were performed intraperitoneally at 8-10 weeks of age using 20 cysts of the ME49 strain of *T. gondii* which were harvested from neural tissue of chronically infected CBA/ca mice. Inhibition of PD-1/PD-L1 signaling was performed by intraperitoneal injection of 1mg of α-PD-L1 (clone: 10F.9G2, BioXcell), while control mice were treated with the IgG2b isotype (clone: LTF-2, BioXcell). Antibody injections were performed one day prior to infection and repeated every 72 hours until indicated for acute or chronic infection studies.

### Assessing Parasite Load and Histology

Serum was isolated posthumously and then analyzed for cytokine concentration by ELISA. 100mg sections of whole liver, heart, thymus, and lungs purified for total DNA using the DNeasy Isolation Kit (Qiagen, MD, USA). DNA concentration was assessed using a Nano Drop spectrophotometer and normalized to equal concentration for qPCR analysis (50ng/ul). qPCR for parasite burden was conducted using toxoplasma specific primers: (forward) 5’-TCCCCTCTGCTGGCGAAAAGT-3’ and (reverse) 5’-AGCGTTCGTGGTCAACTATCGATT G-3’ and Power SYBR Green master mix (Applied Biosystems, CA, USA). The qPCR condition settings were: hold phase (occurs only once at the start): 2min 50C, 10min 95C. PCR phase (occurs 50x): 15s at 95C, 1min @60C. Sections of lung, tissue were fixed in 10% buffered formalin (Jansen Pharmaceuticals, NJ, USA) overnight, then processed for histopathology. Hematoxylin and eosin (H&E) stained sections were used to count for cysts present, or to assess for evidence of leukophilia as result of inhibitory blockade treatment. Cytospins were performed on peritoneal exudate cells (PEC), and H&E stained to allow the quantification of parasites using microscopy.

### Isolation of tissues for analysis

Single cell suspensions were prepared from spleen, lung, liver, bone marrow, and peritoneal exudate cells (PEC) for flow cytometry analysis. Spleens were mechanically processed and passed through a 70μm nylon filter and then lysed in 1ml of 0.846% solution of NH_4_Cl for red blood cell lysis. The cells were then washed and stored on ice. Lungs were harvested and digested with 1mg/ml Collagenase I (Sigma, MO, USA) supplemented with 0.5mg/ml DNAse I (Sigma, MO, USA) in complete RPMI for 45 minutes at 37⁰C. The digested lungs were then passed through a 70μm nylon filter and washed with 10ml of complete RPMI. For liver preparations, the left renal artery was severed, and the mice were perfused using 10ml of 1xDPBS. The gallbladder was removed, and the lobes of the liver were mechanically processed over a 70μm nylon filter and washed. The single cell preparations were then re-suspended in 20ml of 37.5% percoll and centrifuged at 2000rpm for 20mins at RT. The pellet was then re-suspended in NH_4_Cl solution for red blood cell lysis and the cells were then washed and stored on ice. The bone marrow from the femur and tibia of mice was harvested and pooled RBC lysed, and these single cell preparations used for analysis.

### Analysis by flow cytometry

#### T cell staining

Aliquots consisting of 5e6 cells were incubated with Fc block, a mixture of 0.5μg/ml anti-CD16/32 (2.4G2, 553141, BD Biosciences, CA, USA), and 0.25% normal rat serum (10710C, Thermo Fisher Scientific, MA, USA) in a 50μL volume for 30 minutes at 4⁰C. The cells were then washed with ice cold 1x DPBS (21-031-CM, Corning) and stained with Live/Dead Fixable Aqua Dead Cell Stain (L34957, Thermo Fisher Scientific) for 20 minutes on ice and washed in 2% FACS buffer. The cells were then stained with tetramers^60,61^ loaded with the parasite-specific peptides AS15 and Tgd057, and were then washed. The cells were surface stained in a cocktail with the following antibodies: CD4 (GK1.5, 613006, BD Biosciences), α-CD8 (53-6.7, 748535, BD Biosciences), α-CD11a (2D7, 741919, BD Biosciences), α-CD25 (PC61, 102051, Biolegend), α-CD44 (IM7, 103037, Biolegend), α-CD73 (TY/11.8, 127215, Biolegend), α-KLRG1 (2F1, 740279, BD Biosciences), α-GITR (DTA-1, 126315, Biolegend), α-CXCR3 (CXCR3-173, 126531, Biolegend), α-PD-1 (29F.1A12, 135221, Biolegend), α-PD-L1 (10F.9G2, 124319, Biolegend), and α-ICOS (C398.4, 313536, Biolegend) supplemented with brilliant stain buffer (566385, BD Biosciences), for 30 minutes on ice. The cells were washed and re-suspended in Foxp3 Perm-fix cocktail (00-5523-00, Thermo Fisher Scientific) for 4 hours at 4⁰C. The cells were then washed in 1x permeabilization buffer (00-5523-00, Thermo Fisher Scientific) twice, and then re-suspended in an intracellular staining cocktail of: α-CD3 (17A2, 100249, Biolegend, CA, USA), α-Foxp3 (FJK-16s, 11-5773-82, Thermo Fisher Scientific), α-Helios (22F6, 137235, Biolegend), α-Ki67 (B56, 566172, BD Biosciences), α-BCL-2 (BCL/10C4, 633510, Biolegend), α-BIM (K.912.7, MA5-14848, Thermo Fisher Scientific), α-CTLA-4 (UC10-4F10-11, 565778, BD Biosciences), Normal Goat Serum (G9023-10ML, Sigma), and α-Tbet (4B10, 644827, Biolegend) for 2 hours at 4⁰C. The cells were washed with 1x permeabilization buffer twice, and then resuspended with Goat α-Rabbit detection antibody cocktail (A-11046, Thermo Fisher Scientific). The cells were washed and then resuspended in 500μl FACS buffer for flow cytometric analysis.

#### Cytokine staining

To detect intracellular cytokines on T cells, cells were re-suspended in a stimulation cocktail of 0.05ng/ml PMA (Sigma), 0.5ng/ml ionomycin (Sigma), 5ng/ml BFA (Biolegend), Golgi-stop 0.75μl/ml (BD Biosciences, #554724) in cRPMI for 3 hours at 37⁰C and 5% CO_2_. Cells were then washed, and surface stained and permeabilized as described above in the T cell panel. The cytokine stain prepped cells were then intracellularly stained with a cytokine detection panel consisting of: α-IFNγ (XMG1.2, 612769, BD Biosciences), α-IL-10 (JES5-16E3, 505021, Biolegend), α-TNFα (MP6-XT22, 506349, Biolegend), α-Granzyme B (QA16A02, 372208, Biolegend), and α-IL-2 (JES6-5H4, 25-7021-80, Thermo Fisher Scientific) for 2 hours on ice. The cells were washed and then resuspended in 500ul 2% FACS buffer for analysis.

#### Myeloid staining

Splenocytes were washed then stained with mouse α-rat IgG2b (G15-337, 553884, BD Biosciences) for 15 minutes on ice. The cells were washed and live-dead stained and then FC-blocked as described in the T cell panel. The cells were surface stained in an antibody cocktail consisting of: α-B220 (RA2-6B2, 563793, BD Biosciences), α-Ly6G (1A8, 612921, BD Biosciences), α-CD80 (16-10A1, 612773, BD Biosciences), CD45 (30-F11, 748370, BD Biosciences), α-CD3 (145-2C11, 45-0031-82, Thermo Fisher Scientific), α-NK1.1 (PK136, 108728, Biolegend), α-CD11c (N418, 117330, Biolegend), α-Ly6C (HK1.4, 128030, Biolegend), α-CD19 (R35-38, 612978, BD Biosciences), α-CD206 (SA203G11, 150615, Biolegend), α-XCR1 (ZET, 148220, Biolegend), α-MHC-II (M5/114.15.2, 107643, Biolegend), α-CX3CR1 (SA011F11, 149029, Biolegend), α-CD86 (GL1, 17-0862-81, Thermo Fisher Scientific), α-CD11b (M1/70, 101261, Biolegend), α-PD-L1 (MIH5, 12-5982-82, Thermo Fisher Scientific), α-SIRPα (P84, 144016, Biolegend), and α-CD64 (X54-5/7.1, 139314, Biolegend) supplemented with brilliant stain buffer (566385, BD Biosciences) on ice for 30 minutes. The cells were washed and then fixed in FACS buffer supplemented with 2% PFA (15710-S, Electron Microscopy Sciences) for 15 minutes at room temperature. The cells were then washed and then re-suspended in FACS buffer for analysis.

#### Data acquisition

The cells were analyzed on a FACS Symphony A5 (BD Biosciences) using BD FACSDiva v9.0 (BD Biosciences) and analysis was performed with FlowJo (10.5.3, BD biosciences).

### Antigen recall and ELISA

Parasite-specific antigen recall was performed on splenocytes, and the levels of IFNγ were assessed in serum and antigen recall supernatants via ELISA as previously described^62^. α-IFNγ capture antibody (clone AN-18, eBioscience) and biotinylated detection antibody (R4-6A2, eBioscience) were utilized for the ELISA.

### Statistics

Statistical analysis was performed using Prism 8 for Windows (version 8.3) using two-way ANOVA and post-hoc analysis with Sidak corrections for multivariate comparisons. Univariate analysis for individual comparisons were performed with two-tailed student’s *t* test, while univariate analysis with multiple means were assessed using a one-way ANOVA with Tukey tests as post-hoc analysis. Probability for *p* values <0.05 or lower were considered statistically significant. Uniform Manifold Approximation and Projection for Dimension Reduction (uMAP) analysis was performed using the uMAP plug-in (version: 1802.03426, 2018, ©2017, Leland McInness) for Flowjo (Version 10.53). The Euclidean distance function was utilized with a nearest neighbor score of 20, and a minimum distance rating of 0.5. For CD4 T cell, Treg cell, or CD4^+^ Tetramer^+^ T cell uMAP figures, all stained parameters were included in analysis except for: Live Dead (gated out), CD4 (pre-gated), PD-1 (avoiding grouping bias), Foxp3 (avoiding grouping bias or already pre-gated), and in instances of PD-L1 blockade treatment, PD-L1 was excluded from uMAP cluster analysis. The plots generated by uMAP Euclidian algorithms were analyzed via the X-shift tool (version 1.3) to identify unique clusters determined by the uMAP using a weighted k-nearest-neighbor density estimation (kNN-DE) plot. This strategy classifies and defines membership of a cell event into unique cluster groups based on a plurality of observations made from the nearest neighbor events in the uMAP. The X-shift identified clusters were then labeled and subsequently interpreted using the ClusterExplorer (version 1.2.2) tool, which compares X-shift clusters to highlight differences in expression.

## Results

### PD-L1 limits PD-1+ Treg populations at homeostasis

PD-1 is widely expressed by activated CD4^+^ T cell populations and a uMAP analysis of high-dimensional flow cytometry data of bulk splenic CD4^+^ T cells from naïve B6 mice was used to visualize the distribution of this cell surface molecule **(Supplemental Figure 1A-D)**. As expected, few Tconv cells (CD4^+^, Foxp3^−ve^) had high levels of PD-1 whereas approximately 40% of Treg cells (CD4^+^, Foxp3^+ve^) were enriched for PD-1 expression **(Figure 1A)**. Despite exclusion of PD-1 and Foxp3 from the uMAP analytical algorithm, Treg cells naturally segregated from Tconv due to expression of Helios, GITR, and CD25 as well as PD-L1, and CTLA-4 **(Supplemental Figure 1C, 1E)**. The inclusion of additional markers of activation (KLRG1, CD73, and ICOS) highlighted their association with Treg cells, but also illustrated heterogeneity within the Foxp3^+ve^ population at homeostasis **(Supplemental Figure 1F)**. When compared to bulk Foxp3^−ve^ CD4^+^ T cells, Treg cells also have increased levels of activation-associated proteins including CD69, CD11a, CD44, and ICOS, but decreased expression of CD127 **(Supplemental Figure 1G-H)**. To better understand the phenotypic differences between the PD-1^−ve^ and PD-1^+ve^ Treg cells a qualitative uMAP analysis on pre-gated Foxp3^+^ CD4^+^ T cells was utilized that included markers of cellular activation and functional status, but which excluded PD-1 as a calculation factor **(Supplemental Figure 2A-D)**. The clusters were then identified via X-shift analysis and further interpreted using the ClusterExplorer tool **(Supplemental Figure 2E-F)**. Of the 5 subsets that were identified there was a gradation of PD-1 expression (PD-1 ^low^ to PD-1 ^hi^) that overlapped with 4 phenotypic subsets **(Figure 1B-C).** PD-1^−ve^ and PD-1^low^ subsets of the Treg cells were characterized by high expression of CD25 and the pro-survival protein Bcl-2 but were Ki67^low^ **(Figure 1D)**. In contrast, the PD-1^hi^ subset was characterized by increased levels of CD69, and CD11a, which overlapped with the expression of Helios **(Fig 1E)**. Additionally, proteins associated with Treg effector function that included ICOS, CTLA-4, GITR, CD73, PD-L1, and KLRG1 were enriched in the PD-1^hi^ region of the uMAP, **(Fig 1F)**. While not all PD-1^hi^ Treg cells are KLRG1^+^, it is the PD-1^hi^ subset that contains the bulk of effector-associated KLRG1^+^ cells, in addition to ICOS expression and the increased expression of CTLA-4, GITR, CD73, and PD-L1, all while expressing low levels of CD25 **(Fig 1F)**. The heterogeneity of PD-1 expression at homeostasis is illustrated by comparing splenic Treg cells from WT and PD-1 KO mice which allowed the identification of PD-1^−ve^, PD-1^low^, and PD-1^hi^ subsets **(Supplemental Figure 2B)**. Analysis of Treg cells from mice that express nur-77-gfp (a reporter of recent TCR engagement) revealed low Nur-77-gfp expression by conventional CD4^+^ T cells (data not shown) but robust expression by Treg cells associated with expression of PD-1, KLRG1, CD69 and CD11a **(Figure 1G-J)** but loss of CD25 and Bcl-2 **(Figure 1I)**. This combination of low levels of CD25 with increased expression of inhibitory receptors is similar to recently identified eTreg populations^29,33,63,64^ and we will refer to these PD-1^+^ Treg cells as eTreg cells.

**Figure 1:**
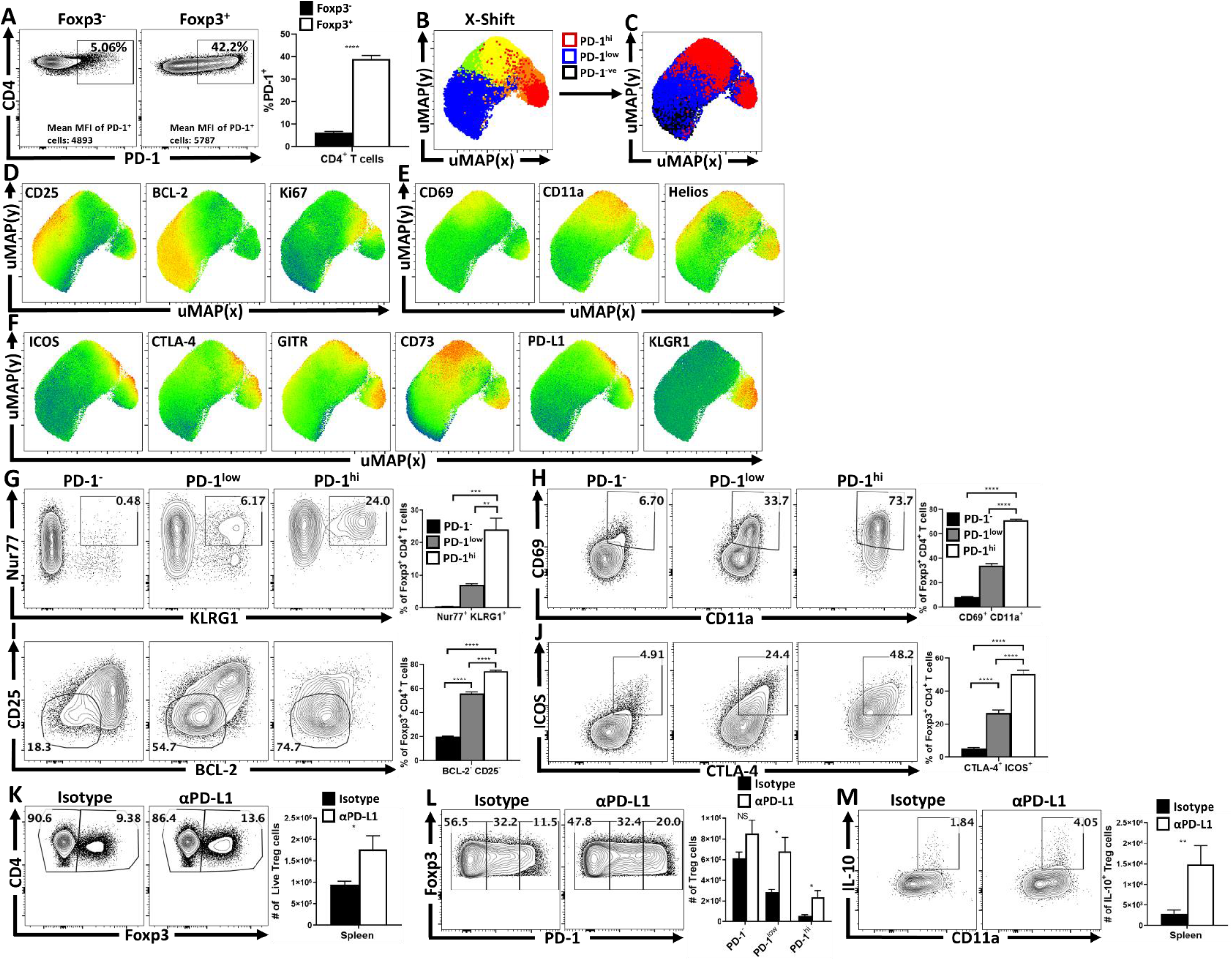
Treg cell heterogeneity at homeostasis and the constitutive impact of Treg cell PD-1. Splenocytes from naïve C57BL/6 mice were analyzed via high-parameter flow cytometry. **(A)** Flow cytometry plots depicting PD-1 expression amongst CD4^+^ Foxp^3−^ T cells (Tconv), in comparison CD4^+^ Foxp^3+^ T cells (Tregs) *(n = 4-5/group student’s t-test, **** = p<0.0001)*. **(B)** uMAP plot of CD4^+^ Foxp^3+^ T cells (Tregs) depicting the X-shift identified subpopulations *(see Supplemental Figure 2 for description)*. **(C)** The same uMAP subdivided into PD-1^−^(black), PD-1^low^ (blue), and PD-1^hi^ (red) regions. **(D)** The median heatmap expression of survival and proliferative-associated proteins CD25, BCL-2, and Ki67. **(E)** The median heatmap expression of activation-associated and nTreg proteins CD69, CD11a, and Helios. **(F)** Heatmap expression of Treg cell effector associated proteins ICOS, CTLA-4, GITR, CD73, PD-L1, and KLRG1. **(G)** Using the PD-1^−^, PD-1^low^, and PD-1^hi^ gating strategy (Supplemental Figure 2), splenocytes from Nur77 reporter mice were analyzed in context of the subdivisions of Treg cells **(G-J)**, comparing the expression of activation and effector proteins Nur77 and KLRG1 **(G)**, CD69 and CD11a **(H)**, CD25 and BCL-2 **(I)**, and ICOS and CTLA-4 **(J)***(n = 5/group, 1-way ANOVA with Tukey multiple comparisons test, **** = p<0.0001)*. Groups of naïve C57BL/6 mice were treated with IgG2b isotype antibody or PD-L1 blocking antibody for 72 hours and their splenocytes were harvested and analyzed via high-parameter flow cytometry. **(K)** Flow cytometry plots of CD4 cells depicting changes in the Foxp^3+^ subset following treatment *(n = 4-5/group student’s t test, * = p<0.05)*. **(L)** Plots of Treg cells depicting an enrichment in the PD-1^low^, and PD-1^hi^ subsets following PD-L1 blockade at homeostasis *(n = 4-5/group, 2-way ANOVA with Sidak’s multiple comparisons test * = p<0.05).* Splenocytes from these hosts were incubated with PMA/ionomycin and stained for cytokine production of IL-10. **(M)** Cytokine staining data depicting an increase in Treg expression of IL-10 following PD-L1 blockade treatment. *(n = 4-5/group student’s t-test, ** = p<0.01)*.

To assess the impact of PD-1/PD-L1 interactions on Treg cells, naïve mice were dosed with an isotype control or αPD-L1 and splenocytes were harvested 72 hours later. Basal levels of PD-L1 were present on multiple immune cell populations and in treated mice αPD-L1 (but not the isotype control) was bound to the surface of these PD-L1+ populations (**Supplemental Figure 3A-B).** This short-term treatment resulted in an increased number of Tregs **(Figure 1K)**, an enrichment in the proportion, and total number of PD-1^+ve^ Treg cells **(Figure 1L)**, their expression of Ki67^+^ **(Supplemental Figure 4E)** and the proportion of and number of PD-1^hi^, CTLA-4^hi^ eTreg cells **(Supplemental Figure 4 D)**. In Nur77-gfp mice, uMAP analysis of the bulk Treg cells (CD3^+^, CD4^+^, Foxp3^+^), highlighted that this blockade increased the proportion of eTreg cells (Nur77^+^, CD11a^hi^, Ki67^+^ which expressed PD-1, CTLA-4 and KLRG1 but were CD25^lo^, **Supplemental Figure 4A-C)**. Congruent with enhanced Treg functions there was an increase in the population of IL-10^+^ Treg cells **(Figure 1M)** and this treatment also resulted in cDC2s and macrophages that expressed decreased levels of CD80 **(Supplemental Figure 4F)**. Essentially similar results were observed with PD-1^−/−^ mice that had an increase in the total number of Treg cells, the number of Ki67^+^ Tregs and an increased population of CD25^low^, BCL-2^low^ eTreg cells (ICOS^+^, CTLA-4^hi^) **(Supplemental Figure 5)**. These findings correlated with increased production of IL-10 and reduced cDC2 expression of CD80 **(Supplemental Figure 5E-F)**. These data sets indicate that at homeostasis, constitutive levels of PD-L1 constrain the size of the PD-1^hi^ eTreg pool.

### Infection–induced IFN-γ promotes PD-L1 expression that limits Treg cells

To determine if PD-1 had a role in the regulation of Treg cell responses during inflammation, WT mice were infected with *T. gondii*, a challenge that results in the collapse of Treg cell populations^47,51^. In naïve mice, PD-L1 was detected on multiple cell types in the spleen and peritoneum, with the highest levels of PD-L1 expression on cDC2s and macrophages **(Figure 2A)**. As early as day 3 post-infection cDC1 and cDC2 increased expression of PD-L1 and there was a marked global upregulation of PD-L1 by neutrophils and monocytes (Figure 2A). This upregulation of PD-L1 was dependent on the early innate production of IFN-γ, as treatment with IFN-γ blocking antibody prevented the upregulation of PD-L1 **(Figure 2A)**. These patterns of expression were maintained at day 10 when bystander CD11a^low^ and activated CD11a^hi^ T cells also expressed significant levels of PD-L1 (data not shown). As expected, by day 10 post-infection there was a marked decrease of the Treg cell populations, however in mice treated with anti-PD-L1 the infection-induced loss of the Treg cells was mitigated **(Figure 2B)**. Further analysis determined that this treatment resulted in the increase of the PD-1^hi^ CTLA4^hi^ eTreg subset **(Figure 2C-D)**. In addition, anti-PD-L1 resulted in an increase in the proportion and number of IL-10^+^ Treg cells **(Figure 2E)**, and a decrease in the proportion and number of CD80^+^ cDC2s **(Figure 2F)**. Surprisingly, the expression of pro-survival BCL-2 on Treg cells was not altered by PD-L1 blockade **(Figure 2G)** but this treatment did antagonize the infection-induced upregulation of the pro-apoptotic molecule BIM **(Figure 2H).**

**Figure 2:**
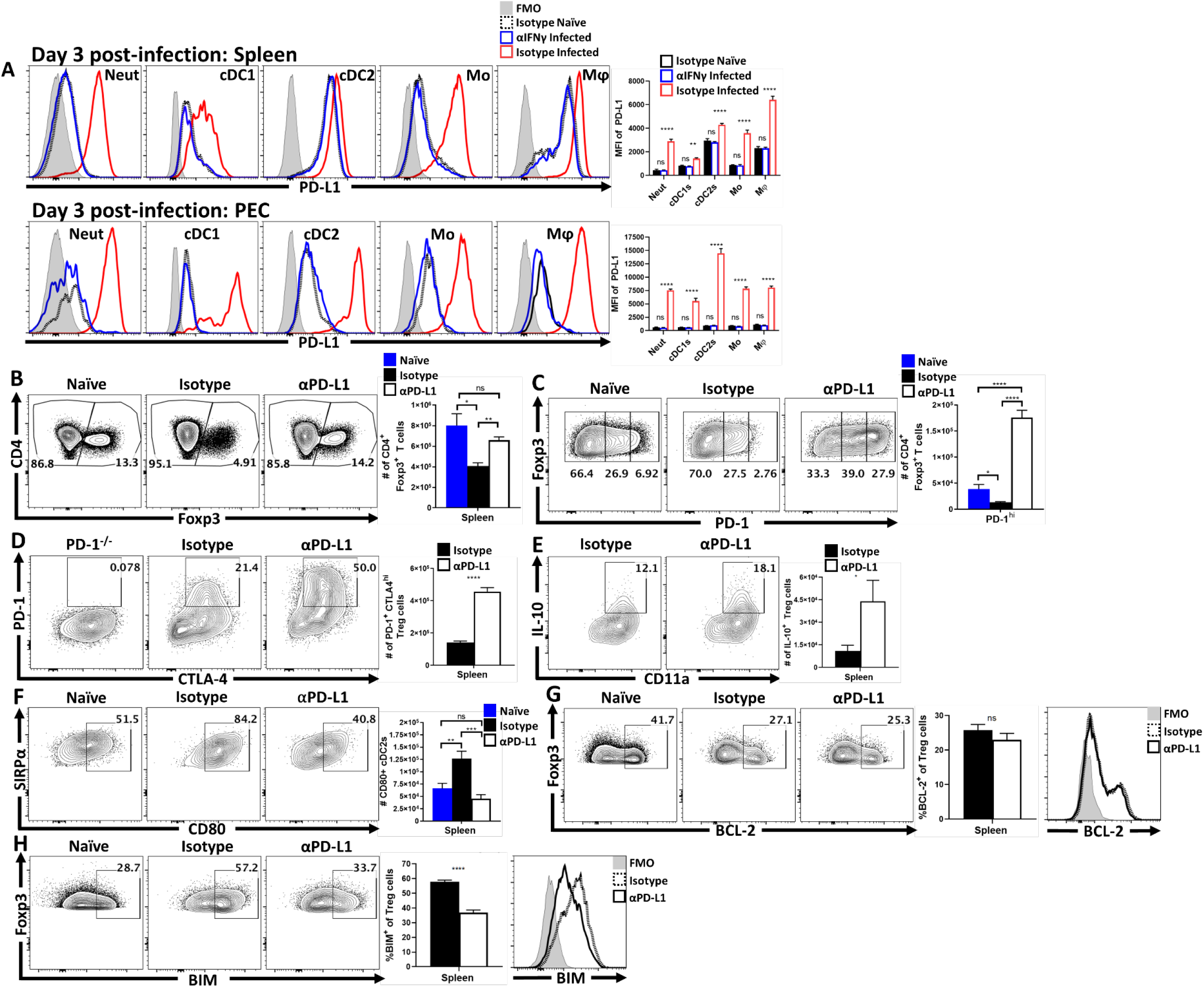
Infection-driven upregulation of PD-L1 drives the crash of PD-1^+^ eTreg cells during the acute phase of infection. Cohorts of C57BL/6 mice were treated with an isotype antibody or IFNγ blocking antibody and half of each group were infected with 20 cysts of ME49 IP. Peritoneal exudate cells (PEC) and splenocytes were isolated 72 hours later and analyzed via flow cytometry. **(A)** Comparative histograms of PD-L1 expression between experimental groups for neutrophils (CD3^−^, B220^−^, CD19^−^, NK1.1^−^, Ly6G^+^, Ly6C^+^, CD11b^+^), cDC1s (CD3^−^, B220^−^, CD19^−^, NK1.1^−^, Ly6G^−^, CD64^−^, CD11c^+^, MHC-II^+^, XCR1^+^), cDC2s (CD3^−^, B220^−^, CD19^−^, NK1.1^−^, Ly6G^−^, CD64^−^, CD11c^+^, MHC-II^+^, SIRPα^+^), monocytes (CD3^−^, B220^−^, CD19^−^, NK1.1^−^, Ly6G^−^, CD64^+^, CD11b^+^, MHC-II^+^, Ly6C^+^), and macrophages (CD3^−^, B220^−^, CD19^−^, NK1.1^−^, Ly6G^−^, CD64^+^, CD11b^+^, MHC-II^+^, Ly6C^−^) *(n = 5/group, 2-way ANOVA with Sidak’s multiple comparisons test ** = p<0.01, **** = p<0.0001)*. Separate groups of mice were then treated with an isotype antibody or PD-L1 blocking antibody and members from each group were infected with 20 cysts of ME49 IP. The antibody treatments were maintained every 72 hours throughout the course of infection until day 10, on which the splenocytes were harvested and analyzed via high parameter flow cytometry. **(B)** Flow plots of bulk CD4^+^ T cells with subsequent gates on the Foxp^3+^ T cells (Treg cells), demonstrating the drop in Treg cells from homeostatic levels during infection, and the maintenance of Tregs during infection with PD-L1 blocking antibody treatment *(n = 5/group, 1-way ANOVA with Tukey’s multiple comparisons test * = p<0.05, ** = p<0.01)*. Flow plots of Treg cells from treatment groups showing enrichment of PD-1^hi^ Treg compartment as a consequence of PD-L1 blockade treatment during infection *(n = 5/group, 1-way ANOVA with Fisher’s test * = p<0.05, **** = p<0.0001)*. **(D)** Flow data for the co-expression of PD-1 and CTLA-4 on bulk Treg cells comparing the number of PD-1^+^ CTLA-4^hi^ Treg cells (eTreg-associated) from animals from the infected groups, using cells from a PD-1^−/−^ host as a gating control *(n = 5/group student’s t-test, **** = p<0.0001)*. **(E)** Splenocytes from each group were stimulated and stained for cytokines, depicted here is IL-10 staining of Treg cells with an in the proportion and total number of IL-10^+^ Treg cells in context of PD-L1 blockade *(n = 5/group student’s t-test, ** = p<0.0001)*. **(F)** Flow plots comparing CD80 expression on cDC2 cells from naïve hosts, and infected hosts treated with isotype or PD-L1 blocking antibody *(n = 5/group, 1-way ANOVA with Tukey’s multiple comparisons test ** = p<0.01, *** = p<0.001)*. **(G)** Flow cytometry analysis of anti-apoptotic BCL-2 expression on bulk Treg cells between the three groups, showing no clear difference in BCL-2 expression when comparing infected groups regardless of anti-PD-L1 treatment *(n = 5/group student’s t-test, ** = p<0.0001)*. **(H)** Comparisons of pro-apoptotic BIM expression on Treg cells, demonstrating a decrease in BIM in infected anti-PD-L1 treated hosts *(n = 5/group student’s t-test, ** = p<0.0001)*.

### Impact of PD-L1 blockade on parasite-specific effector T cells

Infection with *T. gondii* results in reduced production of IL-2 that correlates with the loss of Treg cells^47,51^. Since signals through PD-1 limit effector T cell responses, including production of IL-2^65^, it was possible that systemic blockade of PD-L1 during infection would result in enhanced parasite specific CD4^+^ T cell responses and increased IL-2 that preserve the Treg cell populations. However, while infection resulted in reduced T cell production of IL-2, blockade of PD-L1 did not reverse this effect **(Figure 3A)**. Analysis of the parasite-specific T cell responses at 10 dpi revealed that anti-PD-L1 resulted in a reduction in the magnitude of Tbet^+^ IFNγ^+^ Th_1_ CD4^+^ Tconv cells **(Figure 3B)** and that PD-L1 blockade reduced the number of parasite-specific tetramer^+^ CD11a^hi^, CD4^+^ T cells **(Figure 3C)**. These changes were associated with a reduction of the parasite specific Th_1_ associated Tbet^+^, KLRG1^+^ terminal effector subset **(Figure 3D)**. Notably, the weight loss that accompanies the acute phase of this infection is mediated by the activation of CD4^+^ T cells^66^, and in PD-L1 blockade treated animals there was reduced weight loss and improved physical condition and mobility **(Figure 3E, data not shown)**, despite no significant differences in parasite burden **(Figure 3F)**. The ability to compare infected WT with PD-1^−/−^ mice revealed that the absence of PD-1 resulted in the preservation of Treg cells, an enrichment of the eTreg-associated phenotypes (CD25^low^, BCL-2^low^, ICOS^+^, CTLA-4^hi^), an increase in the number of IL-10^+^ Tregs and a reduction in the proportion of CD80^+^ cDC2s, decreased number of Tbet^+^ KLRG1^+^ parasite specific CD4^+^ T cells, but no significant differences in parasite burden **(Supplemental Figure 6)**. Thus, PD-L1 blockade or loss of PD-1 did not result in increased CD4^+^ T cell effector responses, but rather was associated with enhanced eTreg activity and a reduction in the magnitude of parasite-specific CD4^+^ T cell responses.

**Figure 3:**
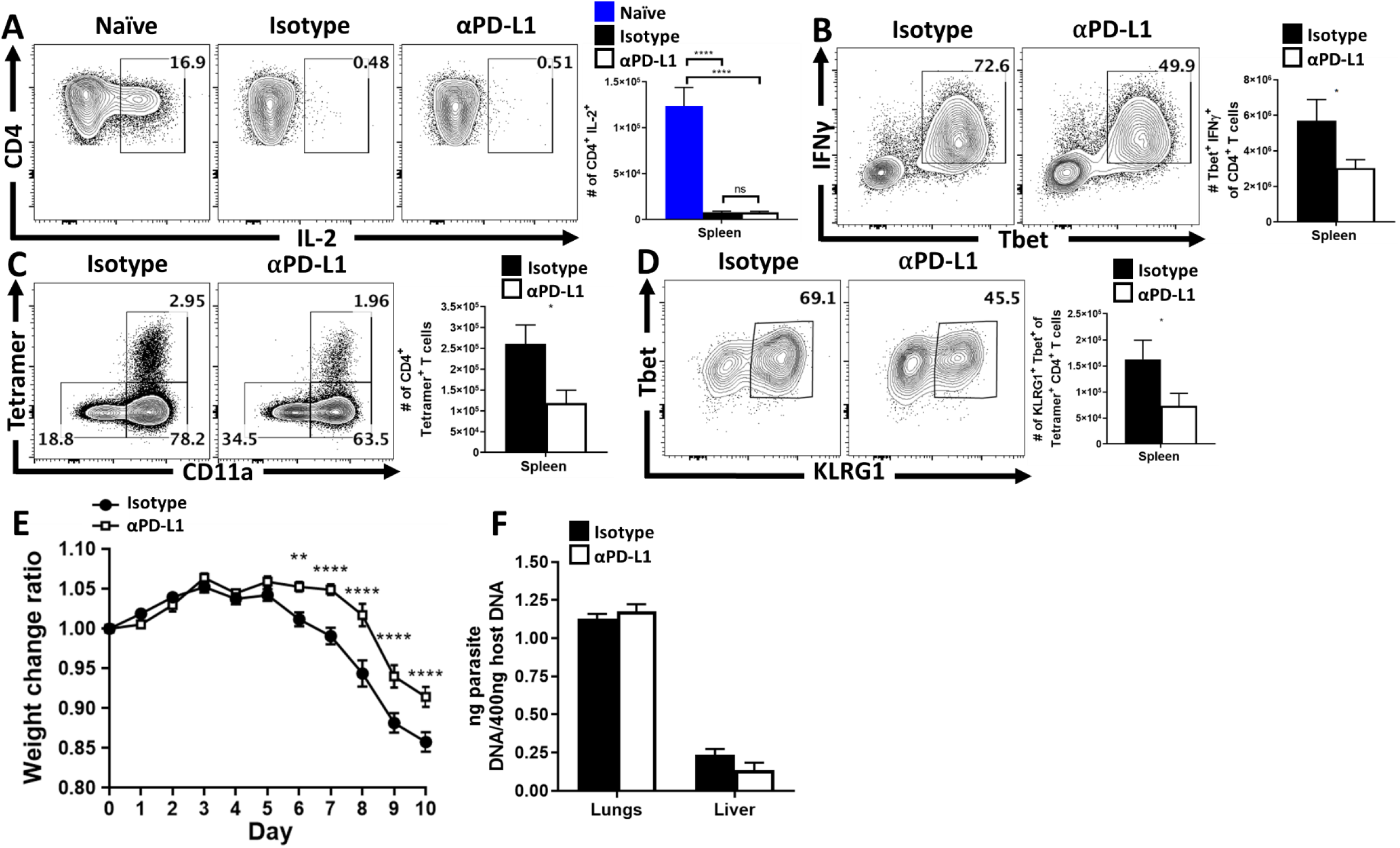
Impact of PD-L1 blockade during infection on effector T cell responses. **(A-B)** Bulk splenocytes harvested from C57BL/6 hosts treated with isotype or PD-L1 blocking antibodies at the tenth day of infection with ME49, were then stimulated with PMA and ionomycin for 3 hours supplemented with brefeldin A and monensin. The cells were permeabilized and then intracellularly stained for cytokines. **(A)** Intracellular stain for IL-2 on bulk Tconv cells (CD3^+^ CD4^+^ Foxp^3−^) from naïve hosts, and infected hosts treated with isotype or PD-L1 blocking antibody *(n = 4-5/group, 1-way ANOVA with Tukey’s multiple comparisons test, **** = p<0.0001)*. **(B)** The magnitude of Th_1_ responses were evaluated via flow cytometry by comparing the proportion and number of IFNγ^+^, Tbet^+^ Tconv cells *(n = 4-5/group student’s t-test, * = p<0.05)*. **(C)** Unstimulated bulk splenocytes were stained using an MHC-II tetramer with the *Toxoplasma*-specific AS15 peptide, comparing the prevalence of parasite-specific CD4^+^ Tconv cells vs CD11a expression between the isotype and anti-PD-L1 treated groups *(n = 5/group student’s t-test, * = p<0.05)*. **(D)** Phenotypic changes to parasite-specific Tconv cells (CD11a^hi^, Tetramer^+^), with a decrease in the proportion and number of Tbet^+^, KLRG1^+^ cells as result of PD-L1 blockade treatment *(n = 5/group student’s t-test, * = p<0.05)*. **(E)** Linear infection timeline graph comparing weight loss ratios between isotype or PD-L1 blockade treatment groups during the acute phase of infection, depicting a reduction in weight loss of PD-L1 blockade treated hosts *(n = 5/group, 2-way ANOVA with Sidak’s multiple comparisons test, ** = p<0.01, **** = p<0.0001)*. **(F)** qPCR analysis of liver and lung samples at day 10 of infection from isotype and anti-PD-L1 treated groups, demonstrating no significant differences observed in parasite burden between control and treated groups despite a decrease in weight loss and Th_1_ Tconv responses. *(n = 5/group, 2-way ANOVA with Sidak’s multiple comparisons test)*.

### Deletion of PD-1 in Treg cells results in increased Treg activities

While the blockade of PD-L 1 or total loss of PD-1 affects eTreg populations it was possible that these effects were mediated indirectly through other immune cell populations. Therefore, to determine if the ability of PD-1 to limit Treg cell responses was intrinsic to Treg cells, Foxp3^Cre^ x PD-1^flox/flox^ mice were generated as recently described^59^. These mice showed normal expression of PD-1 on effector T cell populations while the Treg cell population was PD-1^−ve^ **(Figure 4A)**. At homeostasis, these mice had an increased number of total Treg cells **(Figure 4B)** and while the numbers of the CD25^hi^, BCL-2^hi^ Treg cells appeared normal, the eTreg population was expanded **(Figure 4C-D)**. Thus, PD-1 was not required to generate eTreg cell populations, but expression of PD-1 does limit the size of the eTreg cell pool. In response to infection with *T. gondii*, the Foxp3^Cre^ x PD-1^flox/flox^ mice did not undergo Treg cell collapse **(Figure 4E)** and maintained an enhanced proportion of eTreg cells (defined in the absence of PD-1 as ICOS^+^, CTLA-4^+^, CD25^low^, BCL-2^low^) **(Figure 4F-G)**. Congruent with the previous experiments, there was an increase in the number of IL-10^+^ Treg cells and a reduction in the number of CD80^+^ cDC2s **(Figure 4H-I)**. Experiments that compared mice with a single allele of PD-1 (Foxp3^cre^ x PD-1^wt/flox^) indicated a phenotype intermediate between WT and Foxp3^cre^ x PD-1^flox/flox^ mice **(Supplemental Figure 7)**. Analysis of the CD4^+^ T cell effector responses revealed no evidence of increased IL-2 production **(Figure 5A)**, and a reduced number of parasite specific CD4^+^ T conv cells **(Figure 5B)**. The phenotype of the parasite-specific Tconv compartment was profoundly impacted with a significant decrease in Tbet expression and total number of Tbet^+^ KLRG1^+^ T cells **(Figure 5C-D)**. In contrast to the studies with the PD-L1 blockade or PD-1 KO mice, the lineage specific deletion of PD-1 from Treg cells, resulted in increased weight loss **(Figure 5E)**, and an increase in parasite burden in the liver, lungs, and heart **(Figure 5F)**.

**Figure 4:**
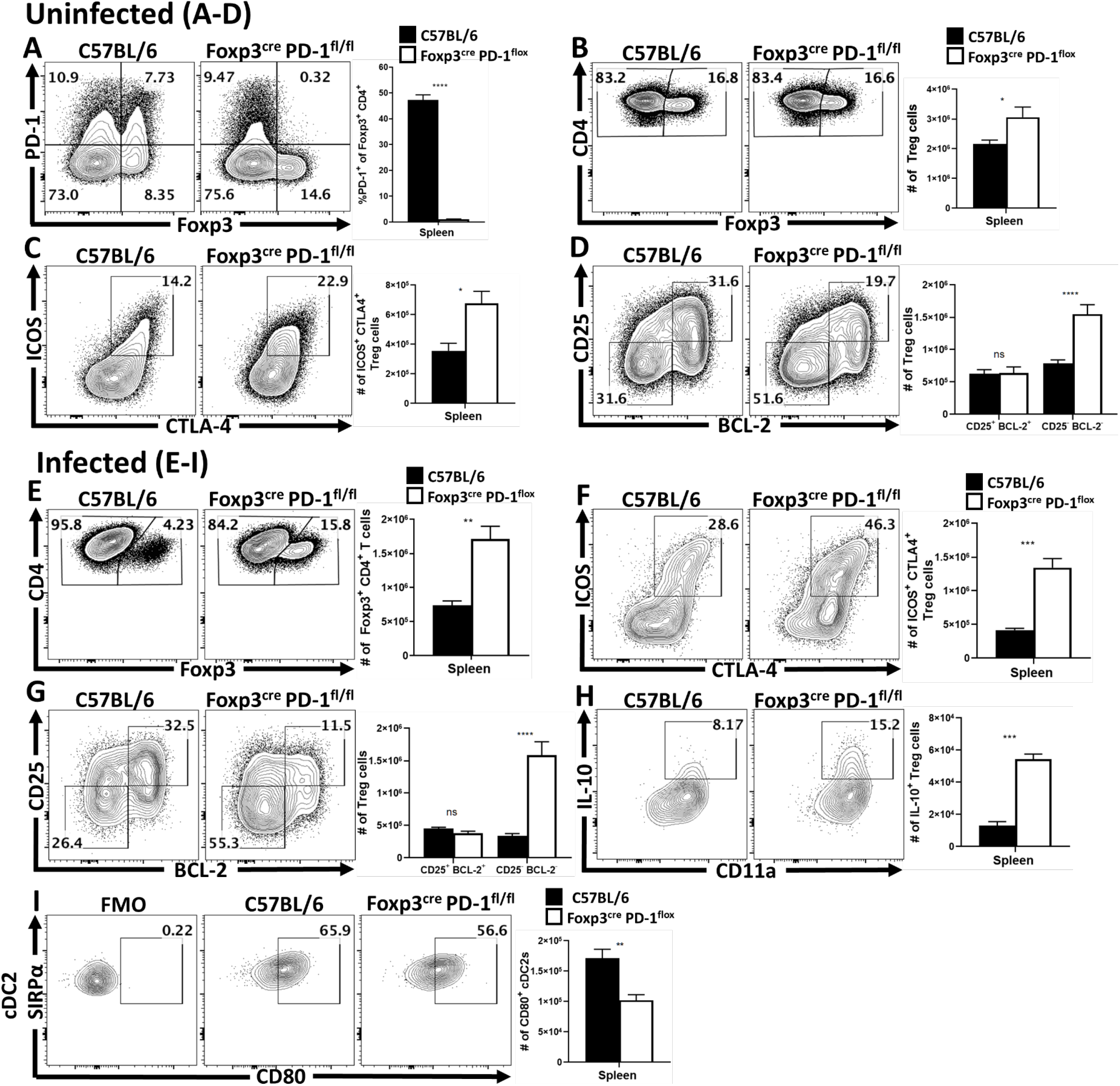
Treg cell-specific deletion of PD-1 enhances eTreg cell populations at homeostasis and prevents Treg cell depletion during infection. **(A-D)** Splenocytes were harvested from naïve C57BL/6 mice and Foxp3^cre^ x PD-1^flox/flox^ mice and analyzed via flow cytometry. **(A)** Bulk CD4^+^ T cells were stained for both Foxp3 and PD-1 to demonstrate a Treg-specific deletion of PD-1, while preserving the expression of PD-1 on the Tconv compartment *(n = 4-5/group student’s t-test, **** = p<0.0001)*. **(B)** Flow cytometry plots of splenic CD4^+^ cells gated on Foxp^3+^ cells (Treg cells), with a result of an increase in the total number of Treg cells in naïve Foxp3^cre^ x PD-1^flox/flox^ mice compared to WT mice at homeostasis *(n = 4-5/group student’s t-test, * = p<0.05)*. **(C)** Treg cell staining of ICOS and CTLA-4, depicting the proportion and number of eTreg-associated (ICOS^+^ CTLA-4^hi^) Treg cells is increased in Foxp3^cre^ x PD-1^flox/flox^ mice *(n = 4-5/group student’s t-test, ** = p<0.01)*, while **(D)** depicts the enhancement is specific to the eTreg compartment (BCL-2^low^, CD25^low^), as the non-eTreg compartment (BCL-2^hi^, CD25^hi^) is consistent in number when compared to WT mice *(n = 4-5/group, 2-way ANOVA with Sidak’s multiple comparisons test, **** = p<0.0001)*. Splenocytes were then harvested from C57BL/6 mice and Foxp3^cre^ x PD-1^flox/flox^ mice at day 10 of infection with ME49 (IP) and analyzed via flow cytometry **(E-I)**. **(E)** Flow plots of splenic CD4^+^ cells gated on Treg cells, demonstrating a preservation of the Treg compartment in cKO mice following infection *(n = 5/group student’s t-test, ** = p<0.01)*. **(F-G)** Pre-gated plots of Treg cells depicting changes in the number of eTreg cells in cKO hosts in context of ICOS^+^ CTLA-4^hi^ *(n = 5/group student’s t-test, *** = p<0.001)*, or BCL-2^low^, CD25^low^ *(n = 5/group, 2-way ANOVA with Sidak’s multiple comparisons test, **** = p<0.0001)* following infection. **(H)** Post-stimulation cytokine stain for IL-10 on bulk Treg cells, demonstrating an increase of IL-10^+^ Treg cells in cKO hosts *(n = 5/group student’s t-test, *** = p<0.001)*, while subsequently **(I)** depicts a reduction in CD80 expression amongst cDC2s in cKO hosts *(n = 5/group student’s t-test, ** = p<0.01)*.

**Figure 5:**
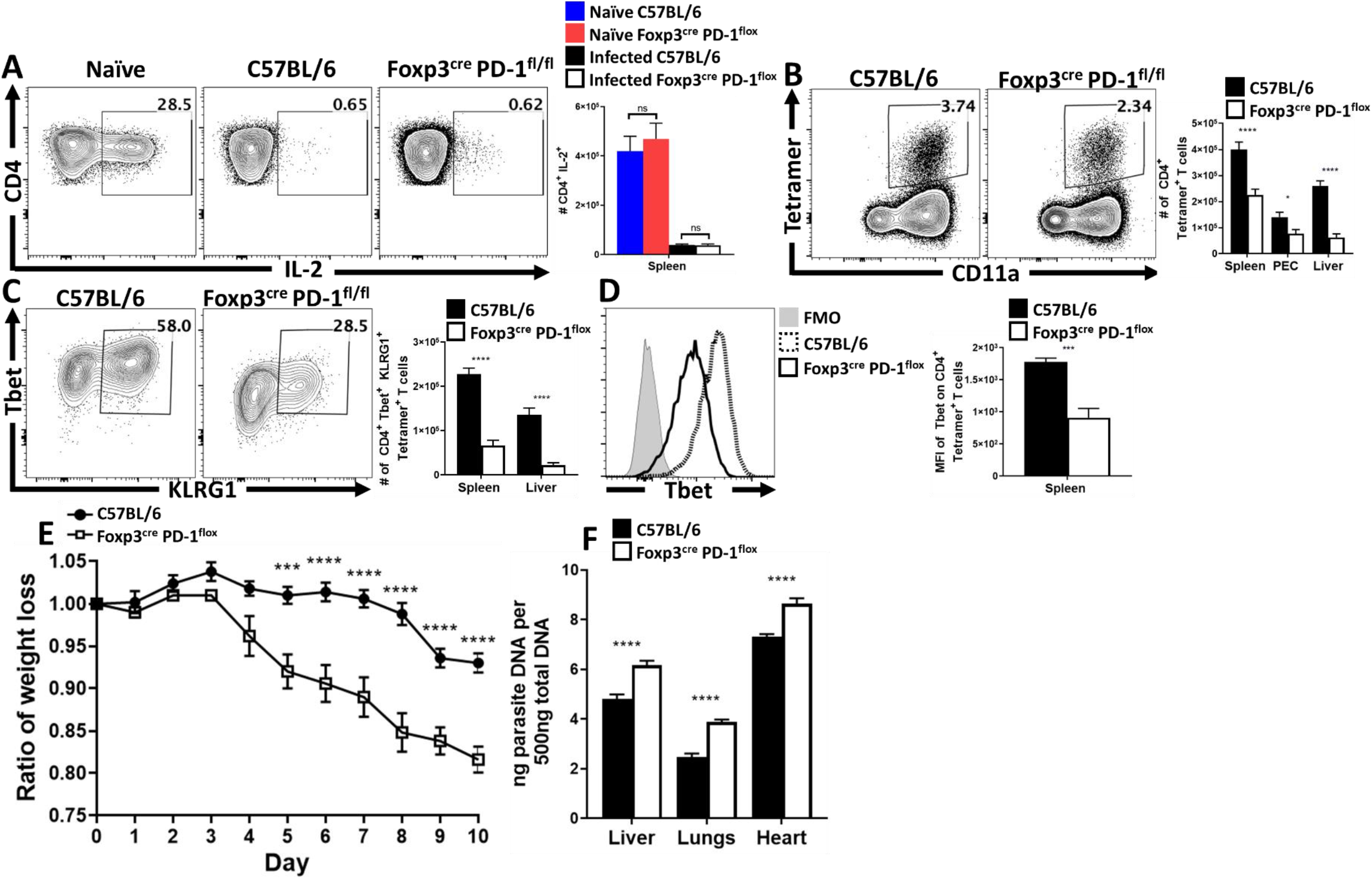
Treg cell-specific deletion of PD-1 results in reduction of parasite specific Th_1_ cells, and a systemic increase in parasite burden. Bulk splenocytes harvested from uninfected and infected groups of mice, consisting of C57BL/6 and Foxp3^cre^ x PD-1^flox/flox^ mice on the tenth day of infection with ME49, were stimulated with PMA and ionomycin for 3 hours supplemented with brefeldin A and monensin. The cells were then permeabilized and intracellularly stained for cytokines. **(A)** Post-stimulation cytokine stain of IL-2 expression on Tconv cells comparing changes between naïve hosts, infected WT and cKO mice, demonstrating no changes to IL-2 production in either uninfected or infected host groups *(n = 4-5/group, 1-way ANOVA with Tukey’s multiple comparisons test, **** = p<0.0001)*. **(B)** Flow cytometry plots of bulk splenocytes from infected WT and cKO hosts, stained for parasite-specific Tconv cells (CD11a^hi^ Tetramer^+^), demonstrating a reduction in parasite specific T cells in cKO hosts across multiple tissues *(n = 5/group, 2-way ANOVA with Sidak’s multiple comparisons test, * = p<0.05, **** = p<0.0001)*. **(C-D)** Parasite specific CD4^+^ T cells from both groups were phenotypically analyzed via flow cytometry for the expression of Tbet and KLRG1, which resulted in a reduction in the raw number of parasite-specific T cells *(n = 5/group, 2-way ANOVA with Sidak’s multiple comparisons test, **** = p<0.0001)* in both the spleen and liver, but also an overall reduction in the MFI of Tbet on these cells *(n = 5/group student’s t-test, *** = p<0.001).* **(E)** Infected cKO hosts demonstrated increased weight-loss throughout the acute phase of infection when compared to WT mice *(n = 5/group, 2-way ANOVA with Sidak’s multiple comparisons test, *** = p<0.001, **** = p<0.0001)*. **(F)** qPCR analysis of parasite burden from liver, lungs, and heart tissue demonstrates an increase in parasite burden in cKO hosts, correlating with a reduction in the number and magnitude of the Th_1_ response. *(n = 5/group, 2-way ANOVA with Sidak’s multiple comparisons test, **** = p<0.0001)*.

In summary, the data demonstrated there is an activated population of PD-1+ Treg cells at homeostasis that are moderated by constitutively expressed PD-L1. Upon primary infection, IFNy mediated inflammation drives the upregulation of PD-L1 which results in the depletion of the suppressive PD-1^+^ eTreg subset. The PD-1/PD-L1 mediated depletion of eTregs subsequently promotes the formation of an effector T cell response to the pathogen **(Figure 6)**.

**Figure 6:**
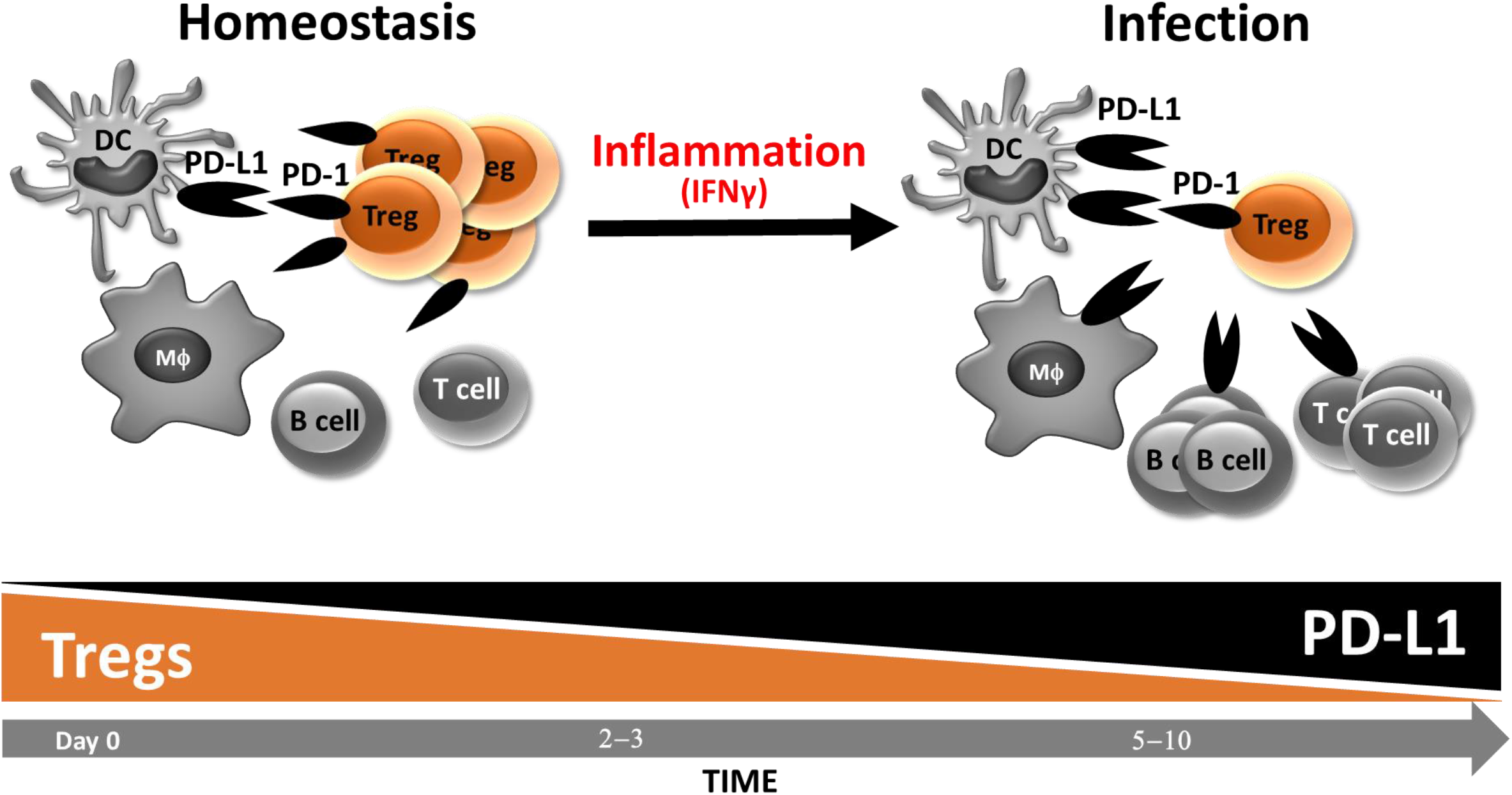
Treg cell PD-1 moderates eTreg cells at homeostasis, and infection-induced inflammation can drive PD-L1 mediated depletion of eTreg cells. At homeostasis, there is an antigen experienced population of PD-1^+^ eTreg cells in addition to PD-L1 expressing APC subsets (cDC2s and macrophages). Following PD-L1 blockade at homeostasis, these eTreg cells quickly respond by demonstrating increases in number and expression of IL-10, suggesting PD-1 functions as a constitutive mechanism of eTreg moderation (Figure 1K-M). In response to primary infection, there is a rapid and ubiquitous upregulation of PD-L1 due to parasite-mediated IFNy expression. The increase in PD-L1 precipitates a depletion of eTreg cells in addition to the expansion of inflammatory T cell populations. As such, eTreg PD-1 functions to restrain eTreg populations at homeostasis, while during infection eTreg PD-1 can promote the formation of an immune response by driving a reduction of immunosuppressive eTreg cells.

## Discussion

The majority of studies on the biology of PD-1 have focused on the impact of this pathway on restraining effector T cell function and less is known about its function on Treg cells. Initial studies concluded that PD-1 promotes iTreg formation and, consistent with the role of PD-1 as a negative regulator of inflammation, PD-1 would enhance this regulatory arm of the immune system^44^. In contrast, the data presented here demonstrate that PD-1 acts to constrain a subset of activated Treg cells at homeostasis, and that during infection the IFN-γ-mediated upregulation of PD-L1 leads to a loss of the eTreg cell population. Support for this conclusion from other systems is provided by the report that PD-L1 blockade enhances the function of PD-1^hi^ Tregs during gastric cancer which leads to hyper-progression and metastasis without any further mutations to the tumor itself^33^. There are also reports that PD-1 checkpoint blockade therapy to enhance effector T cell responses was more effective when combined with strategies that also target Treg cells^67^, and the observation that PD-L1 blockade is also associated with increased Treg cell responses^68^. Similarly, the recent report that specific deletion of PD-1 from Treg cells results in reduced severity of experimental autoimmune encephalomyelitis (EAE)^59^. Thus, in the context of cancer, autoimmunity and infectious disease, PD-1 has a cell-intrinsic role in constraining Treg cell activities. It is notable that CTLA-4 also limits Treg cell responses and that PD-1 and CTLA-4 are frequently co-expressed^69^. Treg cell expression of CTLA-4 has been considered one mechanism that allows Treg cells to suppress accessory cell functions^70^, but Treg-specific deletion of CTLA-4 actually enhances Treg number and immunoregulatory potential^56,71^. This literature highlights that studies on immune checkpoint blockade in the setting of cancer or infectious disease will also need to consider the impact of these interventions on Treg cell populations.

While Treg cells require TCR activation to retain their suppressive phenotype and function^13,72^; recent publications have proposed subdivisions between low and high TCR stimulated subsets of Treg cells^28^. Treg cells with greater levels of TCR activation express minimal CD25^63^, and are the IL-10 secreting, GITR^+^, CTLA-4^hi^ subsets^29^; while Treg cell subsets with less TCR stimulation are CD25^hi^ but low levels of IR^27^. This is of relevance because TCR stimulation promotes PD-1 expression^29,35,73,74^, and the data sets presented here correlate TCR activation with the eTreg subset. The presence of high constitutive level of PD-L1 on MHC-II^hi^ subsets of cDC2s and macrophages suggest that eTreg cells can encounter TCR stimulation and PD-1 signaling simultaneously. Since activation of PD-1 engages immune tyrosine-based inhibitory and switch motifs (ITIM and ITSM)^75–77^ to suppress TCR and co-stimulation mediated signals, this may explain how this pathway continually restrains eTreg cell populations. As such, even short term PD-L1 blockade had an impact on the Treg cell compartment, a finding consistent with reports that concluded that interactions of DC with Treg cells is a continuous process critical for Treg cell homeostasis^72,78,79^. These PD-1^hi^ cells express ICOS paired with low CD25 expression and is reminiscent of the report that DC and ICOS are important in the control of eTreg^63,64^. One implication is that PD-L1 levels reflect global levels of inflammation, and the ability of IFN-γ to increase PD-L1 levels provide a mechanism to contract the PD-1^hi^ eTreg cell population to allow development of the effector responses. In other words, increased PD-L1 levels would act through PD-1 as a rheostat that limits the eTreg cell pool.

PD-L1 blockade administered during in chronic toxoplasmosis has been reported to enhance T cell responses and result in improved parasite control^80^. In contrast, others have found consistent with studies reported here, that PD-1^−/−^ mice have a profound basal defect in the ability to produce IL-12 as result of excess IL-10 production, and consequently are more susceptible to *T. gondii*^81^. During infection, the ubiquitous upregulation of PD-L1, likely in response to IFN-y^82^, did not appear to restrict the development of T cell effector responses^83–85^. Rather, PD-L1 blockade resulted in a reduction in the magnitude of the parasite specific CD4^+^ T cell responses accompanied by reduced immunopathology, despite adequate IFN-γ production from T cells and NK cells to restrict parasite growth. However, the most significant biological impact on this balance between protective and pathological responses was observed with the Foxp3^cre^ x PD-1^fl/fl^ animal models, which expressed the highest levels of IL-10 and were the most susceptible to *T. gondii*. While PD-1KO or PD-L1 blockade treated hosts had an expanded eTreg population, Tconv cells likely benefitted from these strategies from more robust TCR activation due to PD-1 mitigation. As such Foxp3^cre^ x PD-1^fl/fl^ hosts began their infection with an enriched pool of eTreg cells which persisted throughout the infection due to a more precise targeting of the Treg PD-1 pathway, leading to a more robust suppression of the inflammatory response.

Treg cells function to limit a wide variety of immune mediated conditions but pathogens can benefit from their ability to dampen effector responses^49,86^. Consequently, it has been proposed that the transient “crash” of Treg cells during infection is a compromise that allows a balanced T cell response to emerge^47,51^. Acute infection with *T. gondii* (and other pathogens) results in suppressed production of IL-2 and treatment of infected mice with IL-2 complexes that target CD25 mitigate the Treg crash^47,51,56^. The data presented here identify an additional mechanism, distinct from IL-2 deprivation, that targets eTreg cells. It is also relevant to note that the PD-1^hi^ Treg cells are CD25^low^ and ICOS^+^ expressing, which are known to be less dependent on the pro-survival effects of IL-2^63^. The inverted relationship between CD25 expression and the PD-1^hi^ eTreg suggests these are a mature subset of Treg cells that are not dependent on IL-2, and in other studies eTreg cells^63,64,87^ have demonstrated resistances to supplementation or blockade of IL-2^63^. Additionally, it was demonstrated that IL-2 is not required for Treg lineage stability^87^, and it is unclear if the eTreg subdivision also reflects differences in nTreg vs iTreg cells^88,89^, but highlights the heterogeneity of Treg cell populations and the presence of multiple mechanisms, (immune checkpoints included) which are part of the regulatory network at homeostasis and during infection.

---

**Supplemental Figure 1:**
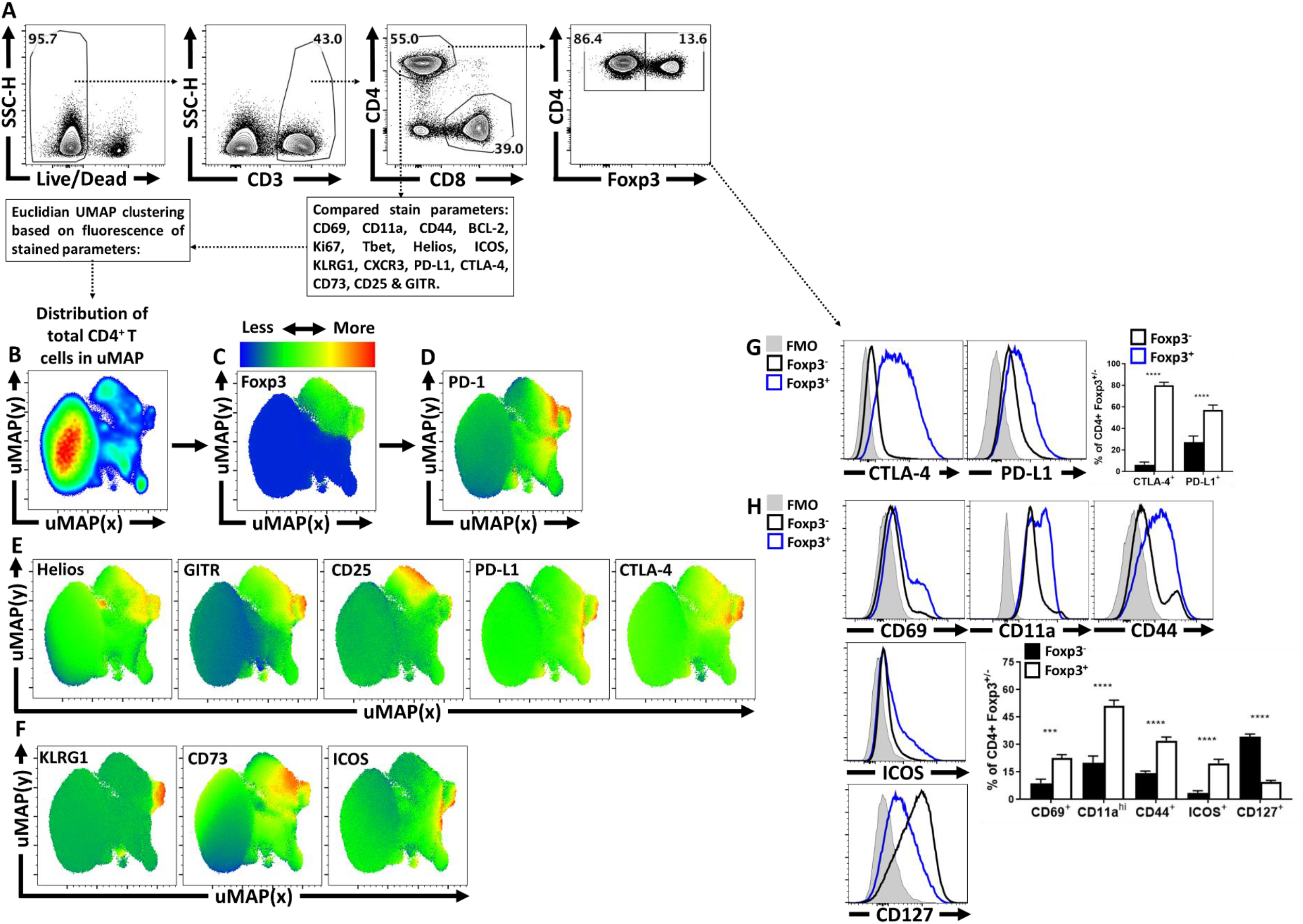
Treg cell heterogeneity at homeostasis and Treg cell expression of PD-1. **(A)** Splenocytes from naïve C57BL/6 mice were analyzed via high-parameter flow cytometry to identify CD4^+^ T cells, and subset them into Foxp^3+^ and Foxp^3−^ subsets. **(B)** Qualitative analysis of bulk CD3^+^, CD4^+^ T cells was conducted to produce a 2-dimensional uMAP representation using dimensional reduction algorithms (excluding CD4, Foxp3, and PD-1 expression as calculated factors). **(C-D)** Regions of CD4^+^ T cells expressing Foxp3 and or PD-1 were identified via median heatmap of expression of the generated uMAP plot. **(E)** The initial distribution uMAP was then qualitatively assessed using median heatmap distribution trends amongst the bulk CD4^+^ T cell pool of Treg cell associated proteins: Helios, GITR, CD25, PD-L1, and CTLA-4, in addition to proteins associated with effector function in Tregs **(F)** KLRG1, CD73, and ICOS. **(G)** Histogram comparisons were then made and quantified between Foxp^3+^ and Foxp^3−^ subsets for the inhibitory proteins CTLA-4 and PD-L1 *(n = 5/group, 2 way ANOVA with Tukey multiple comparisons test, **** = p<0.0001)*. **(H)** Proteins associated with activation (CD69, CD11a, CD44, ICOS, and CD127) were also compared and quantified *(n = 5/group, 2 way ANOVA with Tukey multiple comparisons test, *** = p<0.001, **** = p<0.0001)*.

**Supplemental Figure 2:**
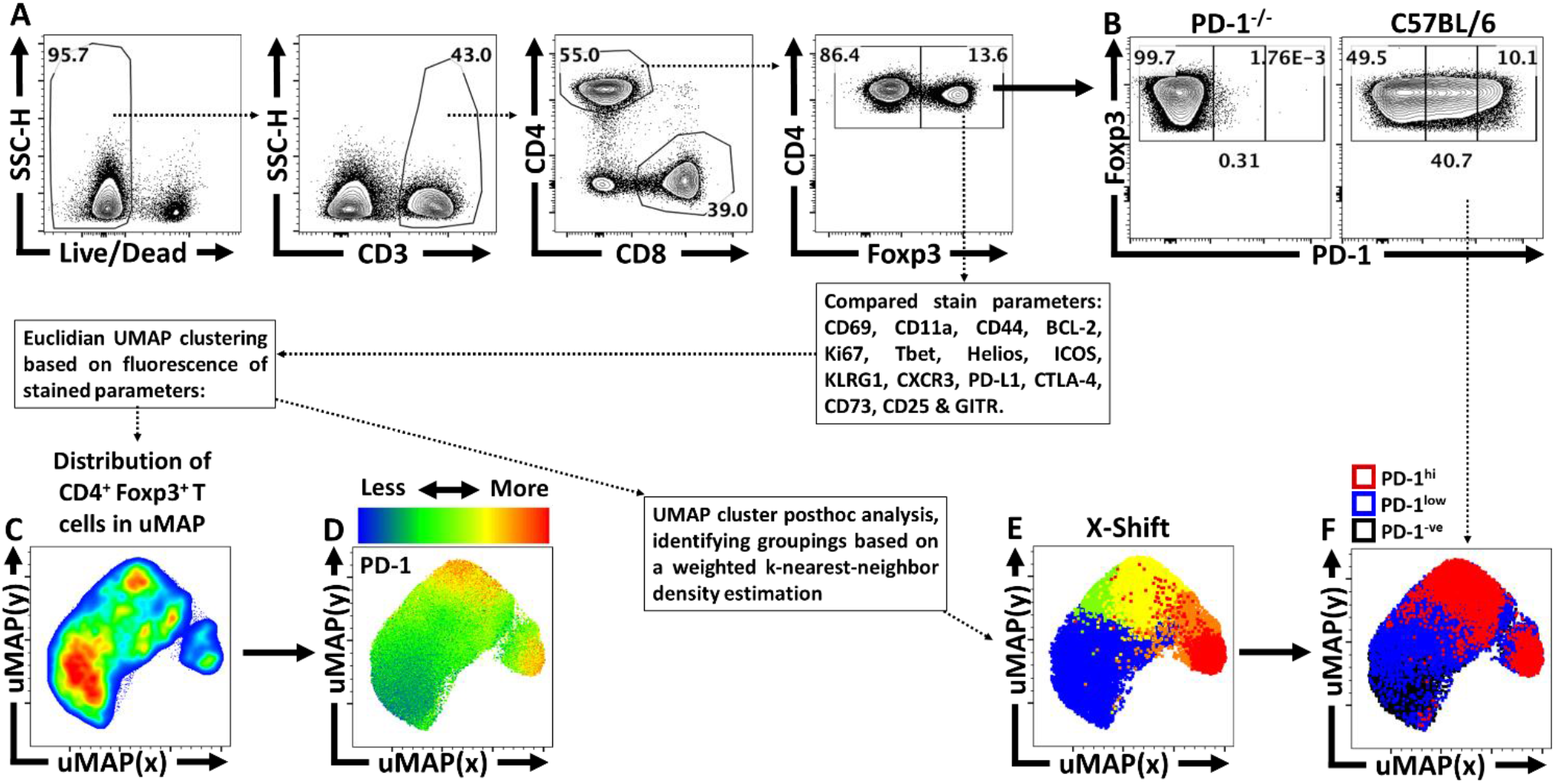
Qualitative X-shift identification of Treg heterogeneity in the PD-1^hi^ cluster of Treg cells. **(A)** Splenocytes from naïve C57BL/6 mice were analyzed via high-parameter flow cytometry to identify CD4^+^ T cells and were then grouped into Foxp^3+ve^ and Foxp^3− ve^ subsets. **(B)** CD4^+^ Foxp^3+^ T cells were then subset into PD-1^−ve^, PD-1^low^, and PD-1^hi^ groups using a PD-1KO host as a negative stain comparative control. **(C)** A uMAP qualitative analysis was generated specifically on CD4^+^ Foxp^3+^ T cells (Treg cells), excluding CD4, PD-1, and Foxp3 as variables in the calculation. **(D)** Depiction of PD-1 expression as a median heatmap amongst the Treg cell uMAP. **(E)** The Treg cell uMAP was then reanalyzed via the X-shift algorithm (excluding CD4, PD-1, and Foxp3 from the calculation) to potentially identify Treg subsets as clusters within the uMAP, with each X-shit identified subset depicted as a separate color. **(F)** Within the same uMAP, the PD-1^−ve^, PD-1^low^, and PD-1^hi^ groups are portrayed as black, blue, and red respectively, to compare the location of these subsets to the locations of the X-shift identified Treg subsets.

**Supplemental Figure 3:**
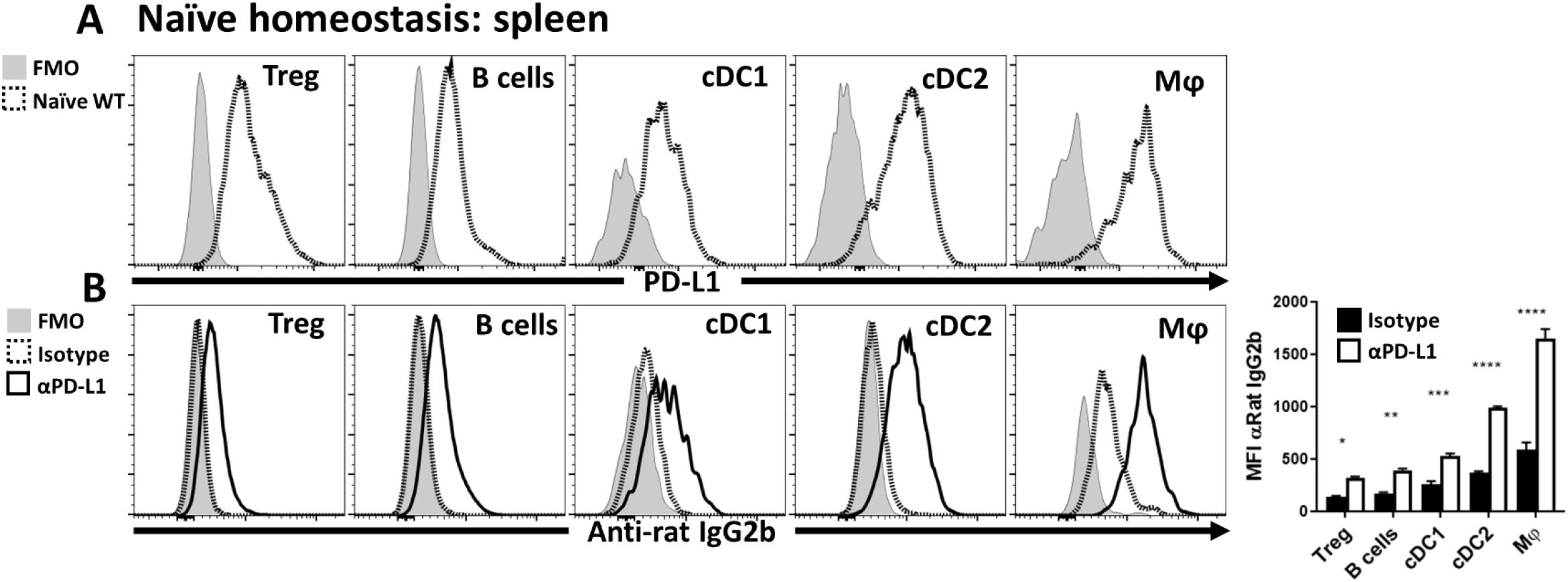
Constitutive PD-L1 expression at homeostasis and anti-PD-L1 blocking antibody detection. **(A)** Splenocytes from C57BL/6 mice were qualitatively analyzed for PD-L1 expression compared to an FMO (fluorescence minus one) via flow cytometry across multiple leukocyte populations: Treg cells (CD3^+^, CD4^+^, Foxp^3+^), B cells (CD3^−^, B220^+^, CD19^+^), cDC1s (CD3^−^, B220^−^, CD19^−^, NK1.1^−^, Ly6G^−^, CD64^−^, CD11c^+^, MHC-II^+^, XCR1^+^), cDC2s (CD3^−^, B220^−^, CD19^−^, NK1.1^−^, Ly6G^−^, CD64^−^, CD11c^+^, MHC-II^+^, SIRPα^+^), and macrophages (CD3^−^, B220^−^, CD19^−^, NK1.1^−^, Ly6G^−^, CD64^+^, CD11b^+^, MHC-II^+^, Ly6Clow). **(B)** C57BL/6 mice were treated with an IP injection of isotype (Rat - IgG2b) or anti-PD-L1 blocking antibody for 72 hours. Splenocytes from these groups were then harvested and stained with an anti-Rat-IgG2b antibody to determine if the PD-L1 blocking antibody was opsonizing the previously identified PD-L1^+^ subsets (Tregs, B cells, cDC1s, cDC2s, and Macrophages). The anti-PD-L1 blocking antibody was readily detected while subsets from the isotype treated animals had minimal anti-Rat-IgG2b staining *(n = 5/group, 2-way ANOVA with Sidak’s multiple comparisons test,* = p<0.05, ** = p<0.01, *** = p<0.001, **** = p<0.0001)*.

**Supplemental Figure 4:**
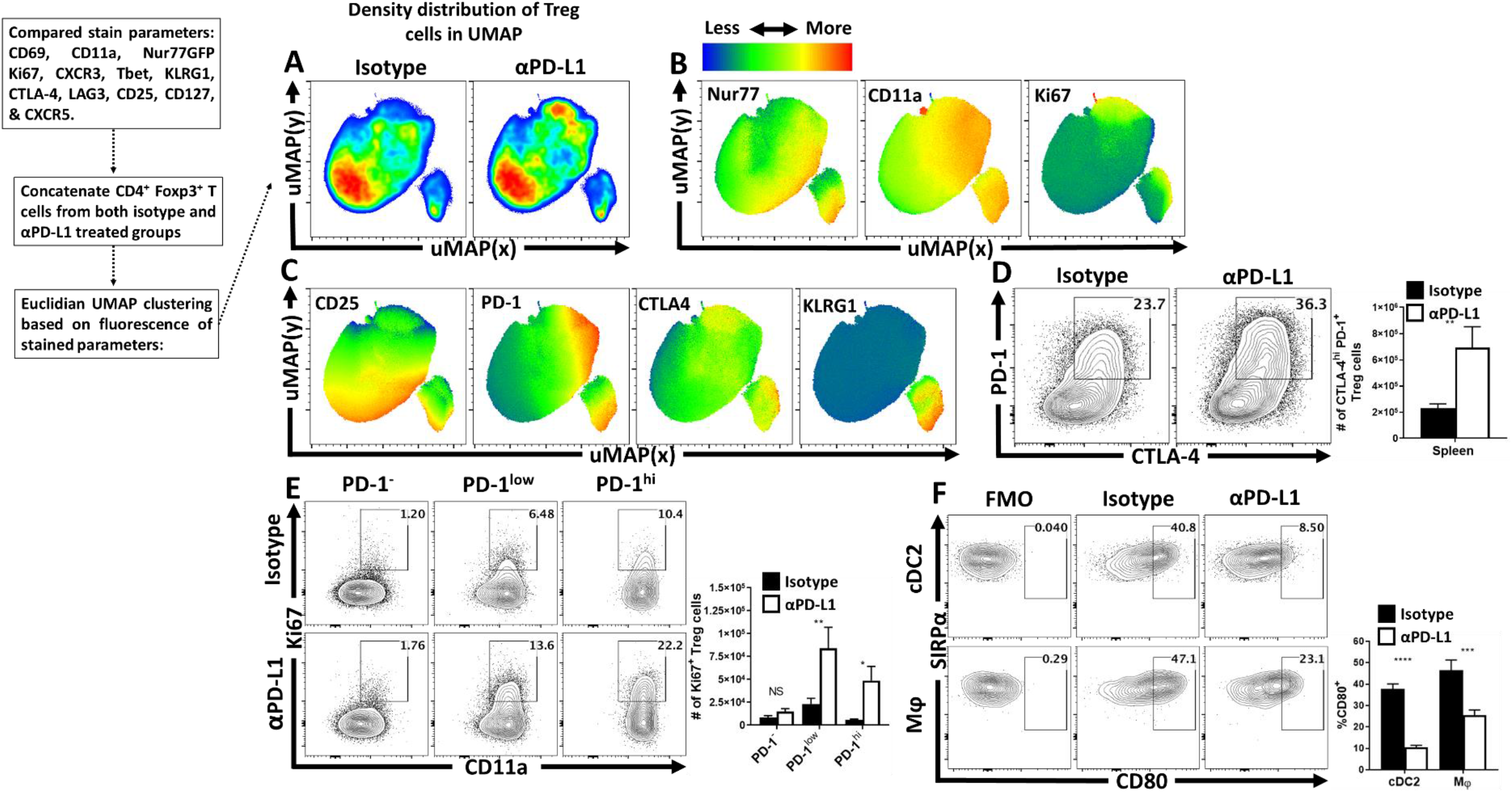
Anti-PD-L1 blockade results in increased eTreg cell activation and proliferation in naïve hosts. **(A-C)** Nur77 reporter mice were treated with a single dose of isotype or anti-PD-L1 blocking antibody for 72 hours. Splenocytes were then harvested and assessed via high-parameter flow cytometry. Treg cell data was then concatenated between the groups and the subsequent qualitative interpretation was conducted via uMAP analysis (excluding Foxp3, PD-1, PD-L1, and CD4 as calculation factors). **(A)** Side-by-side pseudo-color density plot comparison of Treg cells from isotype and anti-PD-L1 treated hosts originating from the same uMAP calculation. There are regional shifts in cell distribution between isotype and anti-PD-L1 treated hosts. **(B)** Heatmap expression analysis across the total combined uMAP data from both groups, depicting median heatmaps of TCR activation associated proteins Nur77, CD11a, and Ki67, with overlapping enrichment of activated Treg cells in anti-PD-L1 treated hosts. **(C)** Additional heatmap analysis of Treg cell associated CD25, inhibitory receptors PD-1 and CTLA-4, and KLRG-1, with an enrichment of overlap between PD-1, CTLA-4, and KLRG1 expression in context of PD-L1 blockade. **(D-F)** C57BL/6 mice were also treated with a single dose of isotype or anti-PD-L1 blocking antibody for 72 hours, and their splenocytes were also isolated and analyzed via high-parameter flow cytometry. **(D)** Flow plot data of splenic Treg cells from isotype and anti-PD-L1 treated hosts comparing changes to the PD-1^+^ CTLA-4^hi^ subset following PD-L1 blockade *(n = 4-5/group student’s t-test, ** = p<0.01)*. **(E)** Treg cells from isotype and PD-L1 blockade treated hosts, gated on activated (CD11a^hi^) cells in cell cycle (Ki67^+^), indicating an increase in PD-1^+^ Treg cells in cell cycle following treatment *(n = 4-5/group, 2-way ANOVA with Sidak’s multiple comparisons test, * = p<0.05, ** = p<0.01)*. **(F)** Flow cytometry data of cDC2s (CD3^−^, B220^−^, CD19^−^, NK1.1^−^, Ly6G^−^, CD64^−^, CD11c^+^, MHC-II^+^, SIRPα^+^), and macrophages (CD3^−^, B220^−^, CD19^−^, NK1.1^−^, Ly6G^−^, CD64^+^, CD11b^+^, MHC-II^+^, Ly6Clow), with gating on CD80^+^ events, depicting a reduction in the proportion of CD80^+^ events following anti-PD-L1 blockade treatment at homeostasis *(n = 4-5/group, 2-way ANOVA with Sidak’s multiple comparisons test, *** = p<0.001, **** = p<0.0001)*.

**Supplemental Figure 5:**
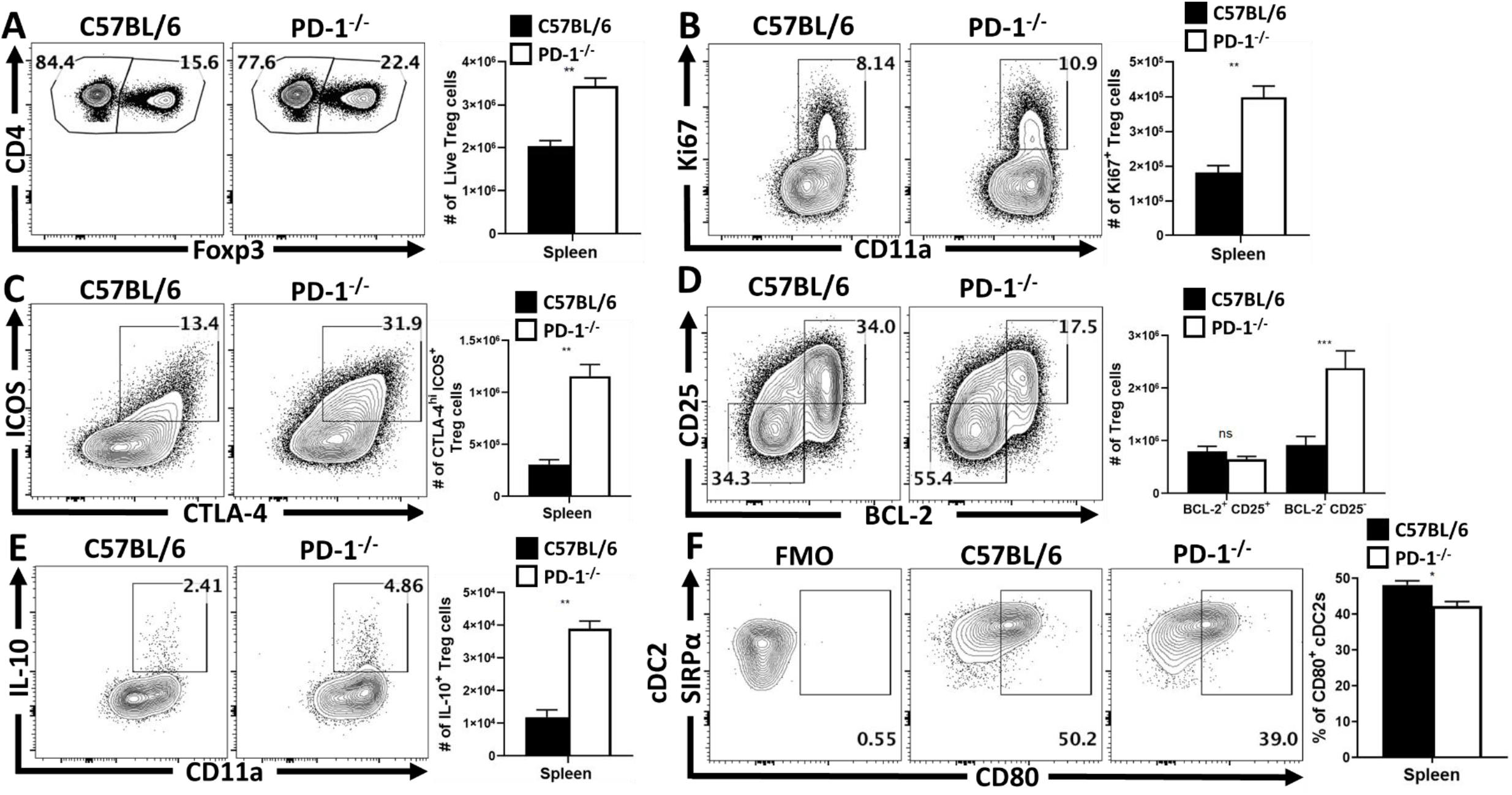
The development of homeostatic eTregs is not dependent on PD-1, and eTregs are limited by PD-1. **(A-F)** Splenocytes from naïve C57BL/6 mice or total PD-1^−/−^ mice were isolated and analyzed via high-parameter flow cytometry. **(A)** Pre-gated CD4^+^ T cells gated on Foxp^3+^ events (Treg cells) depicting an enrichment of Treg cells at homeostasis in PD-1^−/−^ age matched hosts *(n = 5/group student’s t-test, ** = p<0.01)*. Comparative flow plots of Treg cells between WT and PD-1^−/−^ hosts with gating on activated Treg cells in cell cycle (CD11ahi Ki67^+^), demonstrating an increase in Tregs cells undergoing proliferation at homeostasis in PD-1^−/−^ hosts *(n = 5/group student’s t-test, ** = p<0.01)*. **(C)** Treg cell staining of ICOS and CTLA-4, depicting the proportion and number of eTreg-associated (ICOS^+^ CTLA-4^hi^) Treg cells is increased in PD-1^−/−^ mice *(n = 5/group student’s t-test, ** = p<0.01)*, while **(D)** demonstrates this enhancement is specific to the eTreg compartment (BCL-2^low^, CD25^low^), as the non-eTreg compartment (BCL-2^hi^, CD25^hi^) is consistent in number when compared to WT mice *(n = 5/group, 2-way ANOVA with Sidak’s multiple comparisons test, *** = p<0.001)*. Splenocytes from isotype and anti-PD-L1 treated groups were also stimulated and stained for IL-10 and analyzed via flow cytometry. **(E)** Flow plots of Treg cells from WT and PD-1^−/−^ hosts gated on CD11ahi IL-10^+^ events, depicting an increase in the proportion and number of IL-10^+^ Treg cells in PD-1^−/−^ hosts *(n = 5/group student’s t-test, ** = p<0.01)*. **(F)** Splenic cDC2 subsets were identified via flow cytometry (CD3^−^, B220^−^, CD19^−^, NK1.1^−^, Ly6G^−^, CD64^−^, CD11c^+^, MHC-II^+^, SIRPα^+^), and gated on CD80^+^ events based on an FMO. In PD-1^−/−^ hosts, there was observed to be a reduction in the proportion of CD80^+^ cDC2 events compared to WT hosts *(n = 5/group, student’s t test, * = p<0.05)*.

**Supplemental Figure 6:**
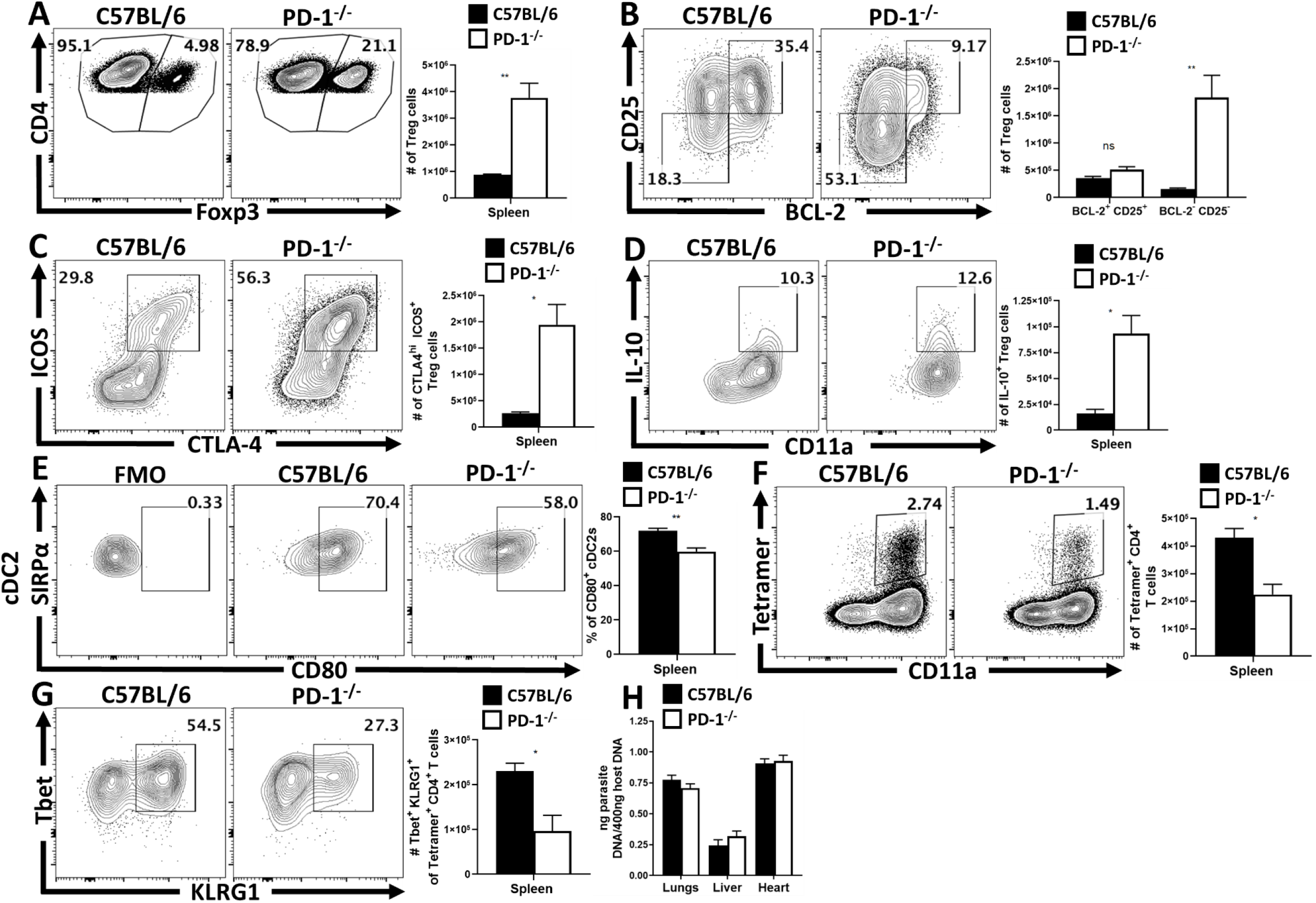
During infection, PD-1^−/−^ mice maintain an increased eTreg pool with diminished parasite specific responses. **(A-G)** C57BL/6 and PD-1^−/−^ mice were IP infected with 20 cysts of *T. gondii* and splenocytes were harvested and analyzed via flow cytometry at day 10 of infection. **(A)** Pre-gated CD4^+^ T cells with gating on Foxp^3+^ events (Treg cells) depicting a preservation of Treg cells in PD-1^−/−^ hosts during infection *(n = 3-5/group student’s t-test, ** = p<0.01)*. **(B)** Treg cell staining of BCL-2 and CD25, demonstrating an eTreg specific increase (BCL-2low, CD25low), as the non-eTreg compartment (BCL-2^hi^, CD25^hi^) is consistent in number when compared to WT mice *(n = 3-5/group, 2-way ANOVA with Sidak’s multiple comparisons test, ** = p<0.01)*. **(C)** Additionally, the proportion and number of eTreg-associated (ICOS^+^ CTLA-4^hi^) Treg cells is increased in infected PD-1^−/−^ mice *(n = 3-5/group student’s t-test, * = p<0.05)*. Splenocytes from WT and PD-1^−/−^ treated groups were stimulated and then stained for IL-10 and analyzed via flow cytometry, **(D)** plots of Treg cells from WT and PD-1^−/−^ hosts gated on CD11ahi IL-10^+^ events, depicting an increase in the proportion and number of IL-10^+^ Treg cells in PD-1^−/−^ hosts *(n = 3-5/group student’s t-test, * = p<0.05)*. **(E)** Splenic cDC2 subsets were identified via flow cytometry (CD3^−^, B220^−^, CD19^−^, NK1.1^−^, Ly6G^−^, CD64^−^, CD11c^+^, MHC-II^+^, SIRPα^+^), and gated on CD80^+^ events based on an FMO. In PD-1^−/−^ hosts, there was observed to be a reduction in the proportion of CD80^+^ cDC2 events compared to WT hosts *(n = 3-5/group, student’s t test, ** = p<0.01)*. **(F)** Splenocytes from infected hosts were tetramer stained using the toxoplasma specific AS15 peptide, and the number of CD11ahi parasite specific CD4^+^ T cells was compared between WT and PD-1^−/−^ hosts *(n = 3-5/group student’s t-test, * = p<0.05)*. **(G)** The phenotype of the parasite specific CD4^+^ T cells (CD11ahi Tetramer^+^) was evaluated for the expression of KLRG1 and Tbet, resulting in a loss of observed Tbet^+^ KLRG1^+^ parasite specific T cells in PD-1^−/−^ hosts *(n = 3-5/group student’s t-test, * = p<0.05)*. **(H)** Parasite burden was assessed via qPCR from tissue samples of lungs, liver, and heart at day 10 of infection, resulting in no significant differences in parasite burden *(n = 3-5/group, 2-way ANOVA with Sidak’s multiple comparisons test)*

**Supplemental Figure 7:**
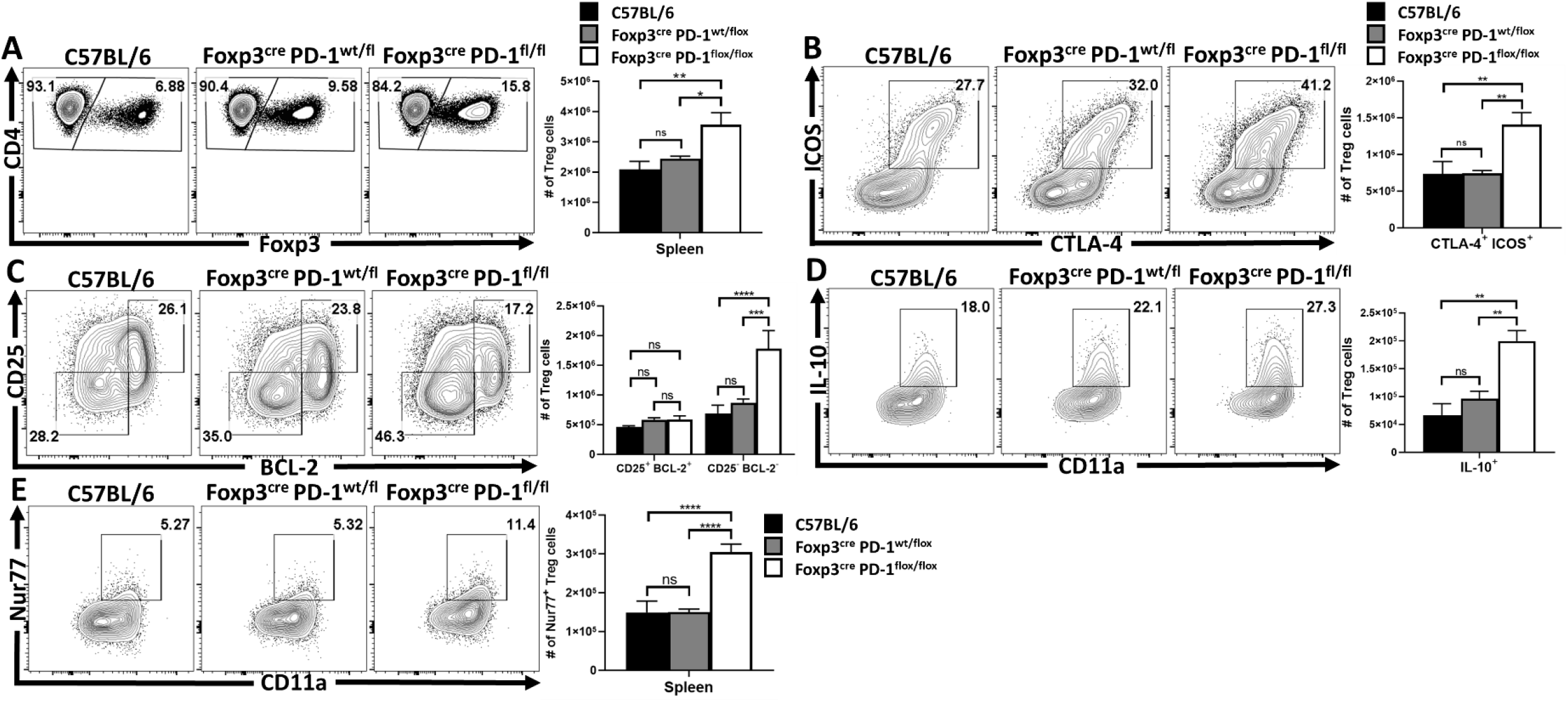
Primary T. gondii infection depletes eTreg cell populations in both C57BL/6 and hemizygous Foxp3^cre^ x PD-1^wt/flox^ mice, while eTreg cells in homozygous Foxp3^cre^ x PD-1^flox/flox^ hosts are spared. **(A-E)** C57BL/6, Foxp3^cre^ x PD-1wt/flox, and Foxp3^cre^ x PD-1^flox/flox^ mice were IP infected with 20 cysts of *T. gondii* (ME49 strain), at day 10 of infection splenocytes were harvested from each group and analyzed via high parameter flow cytometry. **(A)** Flow plots of bulk CD4^+^ T cells from each infected group and were gated on Foxp^3+^ events (Treg cells), depicting similar Treg depletion in WT and hemizygous (PD-1wt/flox) groups, with increased Treg preservation in the homozygous (PD-1^flox/flox^) hosts *(n = 5/group, 1-way ANOVA with Tukey’s multiple comparisons test * = p<0.05, ** = p<0.01)*. **(B)** This increase in Treg cells is likely a result of eTreg survival, as Treg cell staining of ICOS and CTLA-4 between the three groups results in a similar reduction to the proportion and number of eTreg-associated (ICOS^+^ CTLA-4^hi^) Treg cells amongst WT and hemizygous groups, while the homozygous group maintained a significant increase in eTreg associated cells *(n = 5/group, 1-way ANOVA with Tukey’s multiple comparisons test, ** = p<0.01)*. **(C)** As such, since the increase in Treg cells is eTreg associated, there was a specific enhancement to the eTreg associated BCL-2^low^, CD25^low^ compartment in hemizygous mice only, as the non-eTreg compartment (BCL-2^hi^, CD25^hi^) was consistent in number across all three groups *(n = 5/group, 1-way ANOVA with Tukey’s multiple comparisons test *** = p<0.001, **** = p<0.0001)*. Splenocytes from all three groups were also stimulated and stained for IL-10, **(D)** depicts flow plots of Treg cells and their expression of IL-10 vs CD11a. There is no significant change between WT and hemizygous groups, however homozygous mice have a significant increase in the number and proportion of IL-10^+^ Treg cells *(n = 5/group, 1-way ANOVA with Tukey’s multiple comparisons test ** = p<0.01)*. Splenocytes were permeabilized and stained for the downstream TCR-activation protein Nur77 and analyzed via flow cytometry. **(E)** Treg cell plots from the three respective groups depicting no significant differences in Nur77^+^ Treg cells between WT and hemizygous groups, while Treg cells from homozygous hosts have an increased proportion and number of Nur77^+^ Treg cells compared to the other two groups during infection *(n = 5/group, 1-way ANOVA with Tukey’s multiple comparisons test **** = p<0.0001)*.

